# Comprehensive exploration of the translocation, stability and substrate recognition requirements in VIM-2 lactamase

**DOI:** 10.1101/2020.02.19.956706

**Authors:** J. Z. Chen, D.M. Fowler, N. Tokuriki

## Abstract

Metallo-β-lactamases (MBLs) degrade a broad spectrum of β-lactam antibiotics, and are a major disseminating source for multidrug resistant bacteria. Despite many biochemical studies in diverse MBLs, molecular understanding of the roles of residues in the enzyme’s stability and function, and especially substrate specificity, is lacking. Here, we employ deep mutational scanning (DMS) to generate comprehensive single amino acid variant data on a major clinical MBL, VIM-2, by measuring the effect of thousands of VIM-2 mutants on the degradation of three representative classes of β-lactams (ampicillin, cefotaxime, and meropenem) and at two different temperatures (25°C and 37°C). We revealed residues responsible for expression and translocation, and mutations that increase resistance and/or alter substrate specificity. The distribution of specificity-altering mutations unveiled distinct molecular recognition of the three substrates. Moreover, these function-altering mutations are frequently observed among naturally occurring variants, suggesting that the enzymes has continuously evolved to become more potent resistance genes.

## Introduction

The rise of drug-resistant bacterial pathogens has been rapid and inevitable following the introduction of novel antibiotics to the clinic. Pathogens often acquired resistances through horizontal gene transfer (HGT) using mobile genetic elements carrying antibiotic resistance genes, such as plasmids or transposable elements^1–3^. Under constant selection pressure from antibiotic use, these resistance genes continuously evolve to improve their efficacy and alter and broaden their specificity to other classes of antibiotics^2, 4^. Understanding the molecular mechanisms and evolutionary dynamics of antibiotic resistance genes is crucial to finding sustainable solutions against the future dissemination and evolution of antibiotic resistance, such as through predicting future evolution and aiding in antibiotic and inhibitor design^5, 6^.

Metallo-β-lactamases (MBL), or class B β-lactamases, are one of the major sources for the spread of multi-drug resistance bacteria. MBLs are metal dependent hydrolytic enzymes that degrade a broad spectrum of the widely used β-lactam antibiotics, including “last resort” antibiotics such as carbapenems^7^. Plasmid borne MBLs, such as VIM, NDM, IMP, and SPM-types, have been particularly problematic as they can spread to different bacterial pathogens and have no clinically effective inhibitors^8^. All major MBLs have also been undergoing continual evolution; VIM-type MBLs have diversified up to 70 amino acid mutations (26% sequence difference) into over 50 isolated variants, and some variants seem to have developed new substrate specificity^9–13^. Much effort has been made to characterize the molecular mechanisms and identify key residues in several major MBLs using diverse biochemical and structural approaches^14–24^. However, the contributions of the majority of residues in these enzymes remains unexplored, and little is known of the molecular mechanisms governing substrate recognition.

One way to resolve these questions is through comprehensive, large-scale characterizations of mutations affecting the MBLs’ efficacy and specificity. Deep mutational scanning (DMS) is a recently developed method for the characterization of thousands of mutations within a protein using deep sequencing^25–27^. The resulting high-resolution and comprehensive mutational datasets provide invaluable information for deciphering a subset of mutations related to monogenetic disease^28–30^, understanding evolutionary dynamics of proteins—including viral coat, antibiotic resistance genes and hormone receptors^31–34^— as well as elucidating protein sequence-structure-function relationships^35–41^. In particular, conducting DMS on a protein of interest under varying conditions—e.g. against different substrates or in different environments—has further unveiled in-depth molecular details of a protein, such as residues contributing substrate specificity^42, 43^ and protein–environment interactions^44–46^.

In this work, we use DMS to characterize the functional behavior of all ∼5600 single amino acid variants of VIM-2 against three classes of β-lactam antibiotics(ampicillin, cefotaxime, and meropenem) and at two different temperatures (25°C and 37°C), and gain deep insights into the molecular and evolutionary determinants of VIM-2’s behavior. We generate a series of comprehensive and high-quality datasets, and develop a global understanding of VIM-2 by identifying residues that are critical for its function, stability and/or substrate specificity. We also examine VIM-2’s signal peptide—an often overlooked feature despite its importance in expression and transport. Moreover, we use the data to assess their resistance characteristics and rationalize evolutionary outcomes of the clinically isolated natural variants of VIM-type genes, revealing that several mutations in the natural variants are functionally beneficial and lead to changes in substrate specificity.

## Results and Discussion

### Deep mutational scanning of VIM-2 metallo-β-lactamase

DMS was conducted on a library of VIM-2 variants, each encoding a single amino acid substitution. The wild-type (wt) VIM-2 (UniProt ID: A4GRB6) was sub-cloned into a custom pIDR2 vector with a chloramphenicol resistance marker, where VIM-2 expression is controlled by the constitutive ampR promoter (**Supplementary File 1**). We constructed the library of all possible single amino acid variants of wtVIM-2 through PCR based saturation mutagenesis, where each codon position is mutated to an ‘NNN’ codon using restriction-free (RF) cloning (**Fig. 1a**)^47^; there are 5607 possible variants in the library ((20 a.a. + stop codons) × 267 positions). The plasmids of mutagenized codon libraries were pooled into seven groups—six groups of 39 codons (117 bp, 819 variants in each group) and 1 group of 33 (99 bp, 693 variants)—so the mutagenized region of each group can be covered by Illumina NextSeq deep sequencing. We estimated the mutation rate of our library construction by determining the full sequence of 87 variants by Sanger sequencing: only one nucleotide substitution was found outside the intended codon—which corresponds to a mutation rate of 1.4 × 10^-5^—while two other variants had an insertion/deletion. Thus, we constructed a high quality variant library, comparable to other libraries constructed and deep sequenced in a similar manner^42^.

**Figure 1.**
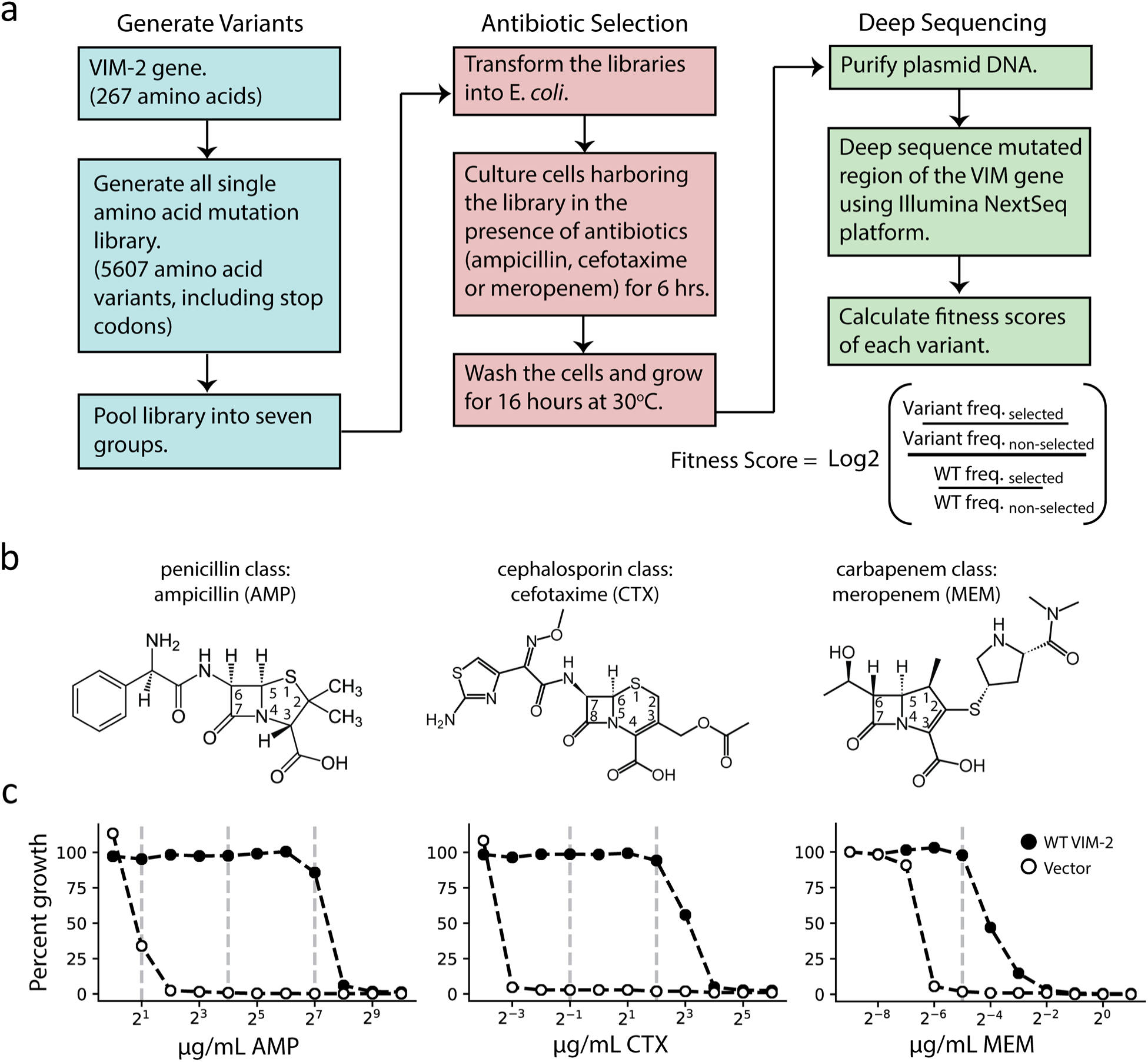
Deep mutational scanning (DMS) overview. (**a**) The workflow for DMS. All single amino acid variants are first generated using RF cloning, subsequently transformed into *E. coli* and then subject to selection for antibiotic resistance conferred to *E. coli*. The effects of selection (fitness score) were evaluated by deep sequencing and comparing the enrichment of each variant with and without selection. (**b**) Chemical structures of the antibiotics used in this study. (**c**) The dose-response growth curve of E. *coli* transformed with wtVIM-2, and an empty vector control for each antibiotic. The vertical dashed lines indicate antibiotic concentrations at which selections were performed in this study.

Cultures of *E. coli* cells transformed with VIM-2 libraries (each group was treated separately) were subjected to antibiotic selection by incubating the culture at 37°C with LB media in the presence (selected) and the absence (non-selected) of three different classes of β-lactam antibiotics—ampicillin (AMP), a 3^rd^ generation penicillin, cefotaxime (CTX), a 3^rd^ generation cephalosporin and meropenem (MEM), a carbapenem (**Fig. 1b**). To determine the selection conditions, the growth of *E. coli* cells harboring the plasmid encoding wtVIM-2 was examined at a range of antibiotic concentrations (1.0-1024 µg/mL AMP, 0.625-64 µg/mL CTX, 0.002-2.0 µg/mL MEM) (**Fig. 1c** and **Supplementary Fig. 1**). We chose to test the highest antibiotic concentration where wtVIM-2 can grow almost 100% relative to growth in media without β-lactam antibiotics, and at successive lower concentrations at 8-fold decrements where the range permits; selected conditions are 128, 16 and 2.0 µg/mL of AMP, 4.0 and 0.5 µg/mL CTX, and 0.031 µg/mL MEM (**Fig. 1c**). The selection process for each antibiotic was conducted in duplicate on separate days. After selection, the plasmids were isolated, the mutagenized region of each group was amplified by PCR, and the amplicons were sequenced by the Illumina NextSeq 550 platform. The sequencing reads were error filtered, and the fitness score of each variant relative to wtVIM-2 was calculated using equation (1). (see methods for “deep sequencing and quality control”).

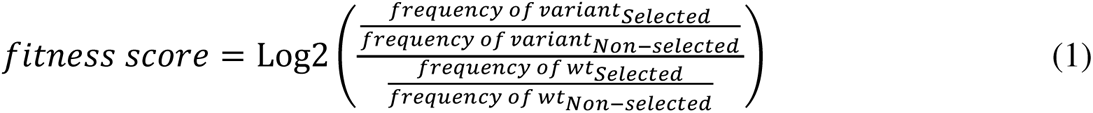

Where the frequency of a variant (or wt) is the deep sequencing read count of the variant divided by the total reads in the corresponding sample. Variants with frequencies below the threshold of deep sequencing errors that was estimated from the deep sequencing of wtVIM2 (see methods for “Variant identification and noise filtering”) were excluded during scoring. The non-selected library shows excellent coverage, with 5535 of 5607 (98.7%) variants present after filtering in at least one replicate while 97.8% are observed in both replicates (**Supplementary Table 1**). For selected libraries, we calculate the fitness score for any variants present in at least one non-selected replicate then average the fitness scores between the two selection replicates (see **Supplementary Data 1** for all fitness scores).

Our DMS experiments show high replicability in all conditions tested. The R^2^ of a linear regression between variants observed in both replicates is 0.94 for selection with 128 µg/mL AMP, 0.91 for 4.0 µg/mL CTX and 0.85 for 0.031 µg/mL MEM (**Fig. 2** and **Supplementary Fig. 2**). As expected, variants with synonymous mutations have near neutral fitness and variants with nonsense mutations (stop codons) have the lowest fitness scores. At the highest concentration of each antibiotic, variants with stop codons have fitness scores centered around −4 and lower, thus a fitness score of −4 is considered the lowest score cut-off for downstream analyses (**Fig. 2**). Like previous DMS studies with other proteins, the overall fitness distribution of all variants exhibit a bi-modal distribution with a peak at neutral fitness and a long tail stretching toward negative fitness to another peak at the cutoff of −4 (**Fig. 2**)^32, 36, 39, 48^.

**Figure 2.**
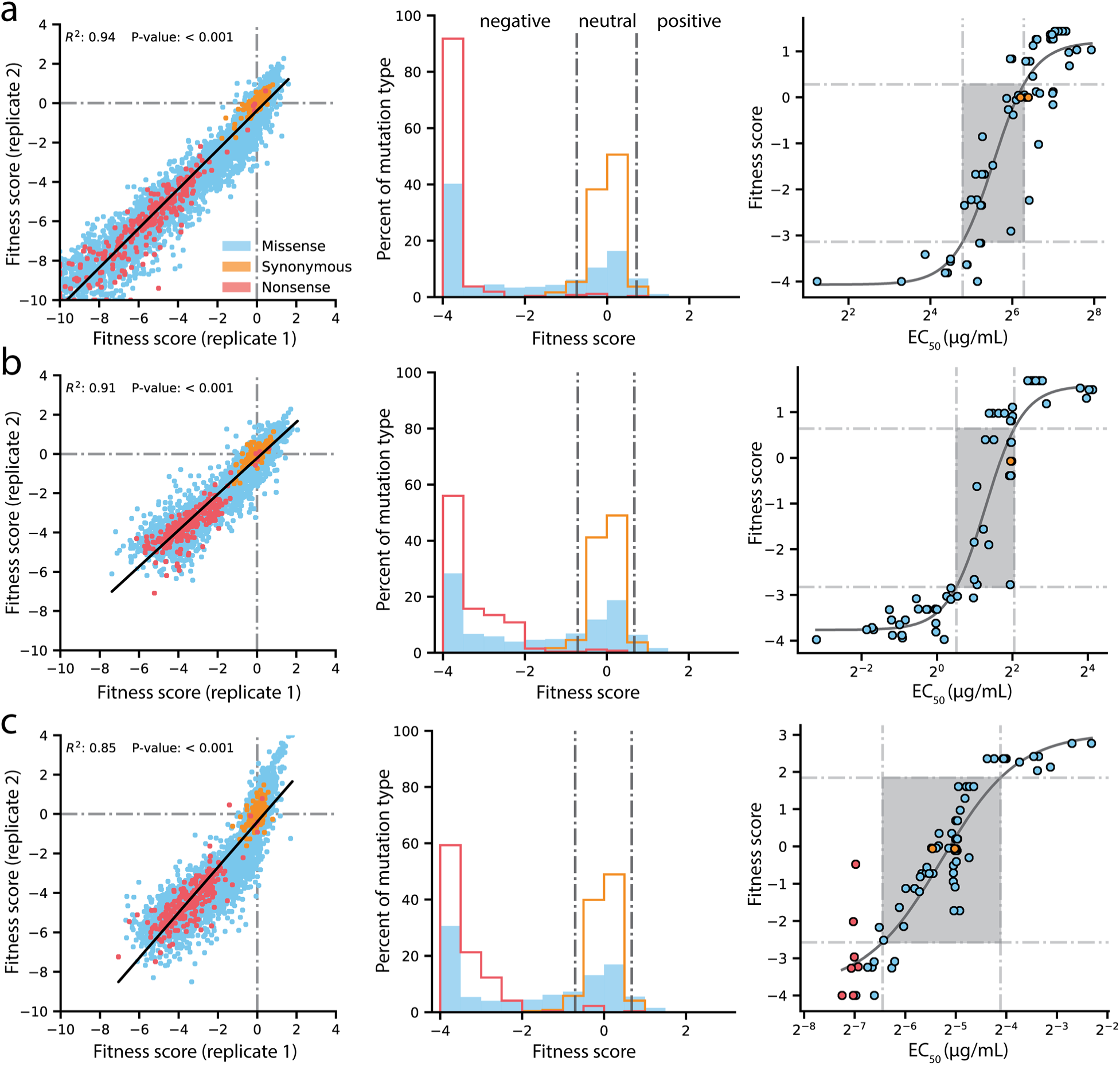
Quality control of DMS data and general mutational properties of VIM-2. In the horizontal panels, data is shown for (**a**) 128 µg/mL AMP selection, (**b**) 4.0 µg/mL CTX selection and (**c**) 0.031 µg/mL MEM selection. For each panel, the left plot shows correlation between fitness scores of all variants in the two replicates of DMS. The middle plot shows distribution of fitness effects for all variants separated into synonymous, missense and nonsense distributions, where the vertical grey lines indicates fitness score cut-offs used to classify fitness effects as positive, neutral or negative. The right plot shows the relationship of DMS fitness scores with antibiotic resistance (*EC_50_*) of isolated variants measured in a dose-response curve; variants with resistance lower than the tested range could not be fitted for *EC_50_*, leading to 69 individually measured *EC_50_* values for AMP, 67 for CTX and 75 for MEM—some points are the same codon or amino acid variant isolated multiple times. The filled rectangle in the background indicates the region of linear association between fitness scores and *EC*_50_. The legend in panel **a**) is shared by all panels.

We confirmed the DMS fitness scores reflect the actual resistance level of variants (**Fig. 2** and **Supplementary Data 2**). We isolated 45 unique variants (61 unique codons), and determined the half maximal effective concentration (*EC*_50_) of *E. coli* culture harboring each variant for three antibiotics by measuring antibiotic dose-response curves. We fit the relationship using a sigmoidal function and identify a linear range of correlation for fitness scores within −3.1 to 0.3 for AMP (*EC_50_* 27–78µg/mL), −2.8 to 0.6 for CTX (*EC_50_* 1.4–4.1µg/mL) and −2.6 to 1.8 for MEM (*EC_50_* 0.011–0.058µg/mL), which correspond to a 2.8, 2.9 and 5.0-fold range of *EC_50_* values for AMP, CTX and MEM, respectively^49^. All fitness scores outside the linear range are still qualitatively consistent with *EC_50_*, where higher fitness scores correspond to higher *EC_50_* values and lower fitness scores correspond to lower *EC_50_* values.

### Global view of VIM-2 enzyme characteristics

The fitness scores for variants selected at 128 µg/mL AMP are shown in **Fig. 3** (See **Supplementary Fig. 3-4** for CTX and MEM). At a glance, there are several interesting trends in the DMS data of VIM-2. Variants with Cys mutations are highly deleterious throughout the catalytic domain (positions 27-266). As wtVIM-2 possesses only one Cys for metal binding, additional Cys may cause the formation of undesired disulfide bonds, leading to misfolding. Pro variants are also often deleterious, as this residue disrupts secondary structures^32, 36, 41^. We found 112 positions (42% of all residues) are highly sensitive to mutations, where over 75% of missense variants (excluding synonymous and nonsense mutations) display a fitness score < −2.0. These positions are likely key requirements for catalytic activity or protein stability and folding in the wtVIM-2. Indeed, these positions include all six active-site metal coordinating residues (His114, His116, Asp118, His179, Cys198, His240), as well as 3-4 residues adjacent to each metal binding residue in the amino acid sequence that are likely to play important roles in the metal configuration and enzymatic function. Additionally, 92% of positions with high mutational sensitivity— including all metal binding residues—are located in the core of the protein (accessible surface area, ASA, of residue < 30%) and 63% are almost completely buried (ASA < 5%), congruent with previous findings^25, 32, 37, 42, 46, 50^ (**Fig. 4a**).

**Figure 3.**
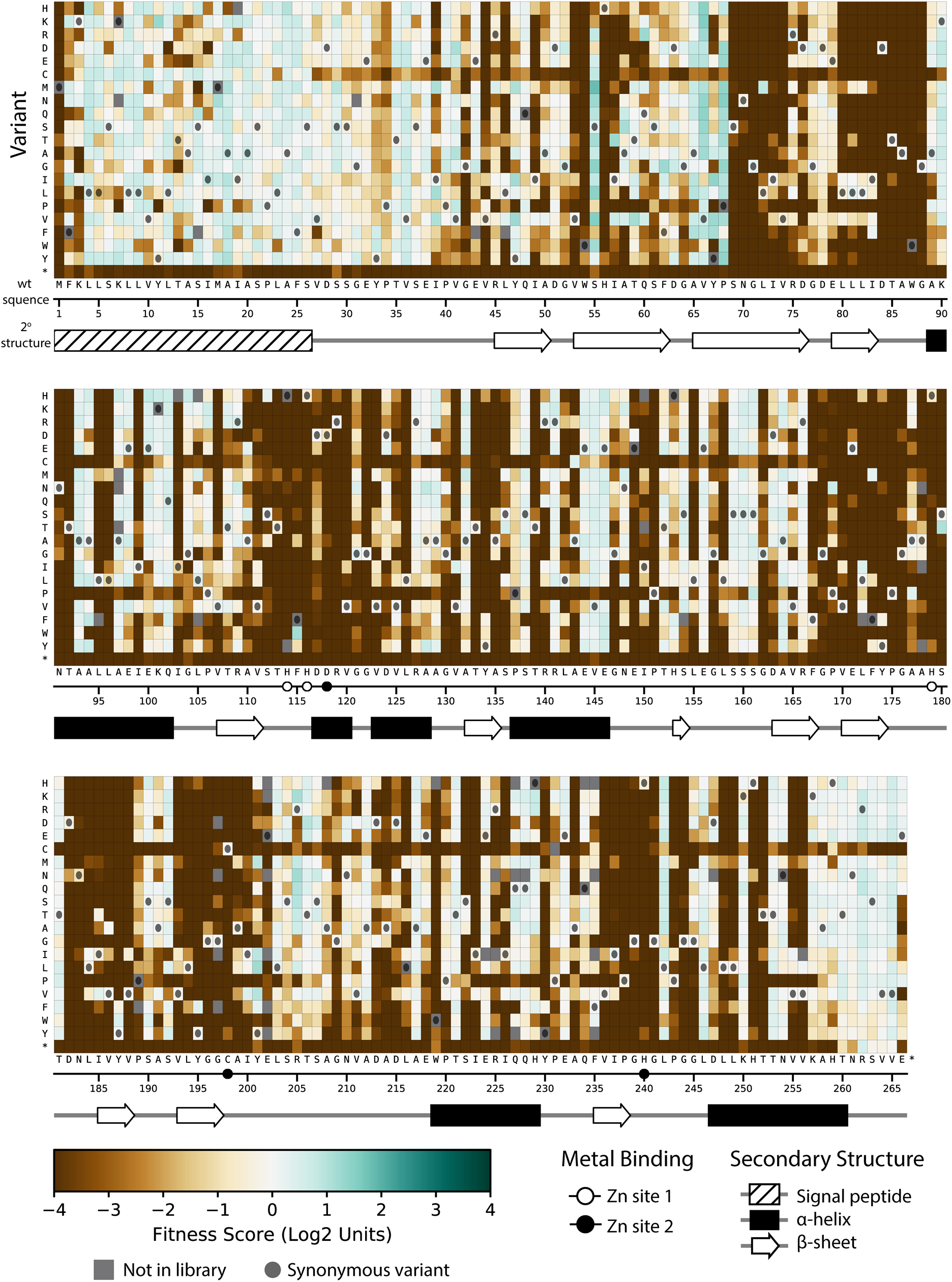
Fitness of all VIM-2 single amino acid variants under 128 µg/mL AMP selection. Each cell in the heat map represents the fitness score of a single amino acid variant. Synonymous variants are indicated by black circles and variants that are not present in the library are in grey. The x-axis under the heatmap indicates the wt residue and position (the 6 active site metal binding residues are highlighted as circles), while the y-axis indicates the variant residue at that position. The secondary structure of the wtVIM-2 crystal structure (PDB: 4bz3) is displayed below the heatmap.

**Figure 4.**
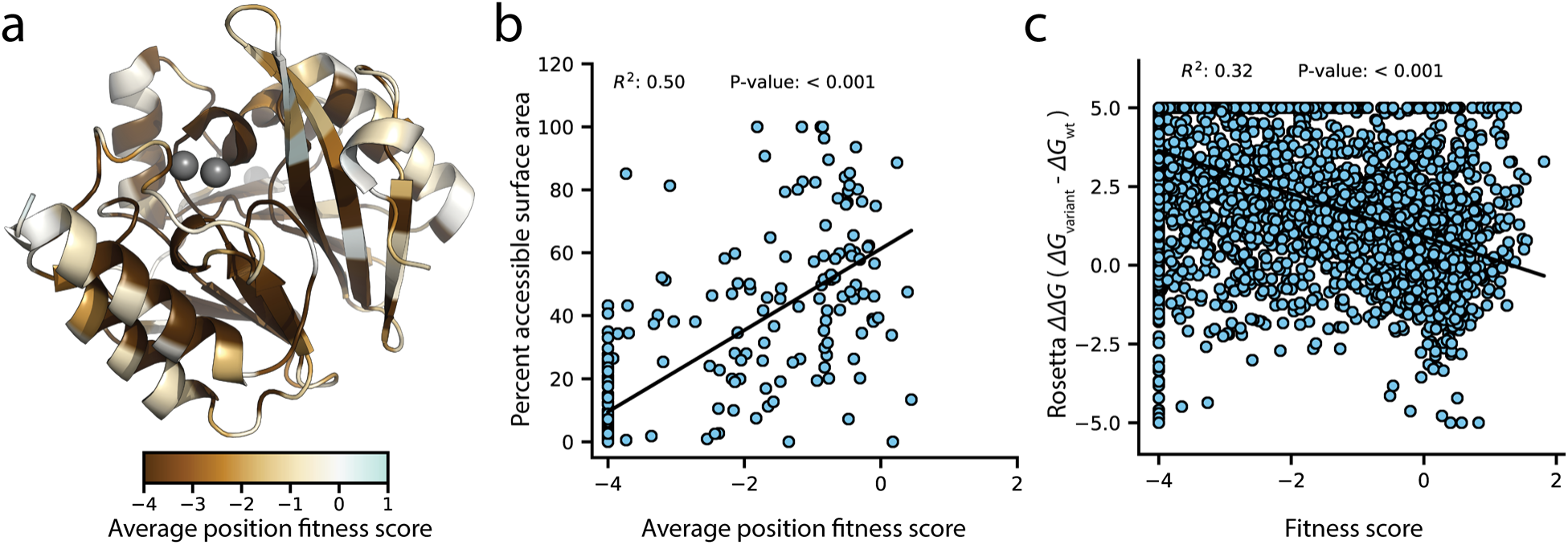
Correlation of fitness with structural attributes. Fitness scores are from DMS at 128 µg/mL AMP selection. (**a**) Crystal structure of wtVIM-2 (PDB: 4bz3) colored by the average fitness of 20 amino acid mutations at each position. (**b**) The correlation between accessible surface area and the average fitness of 20 amino acid mutations at each position. (**c**) The correlation between the changes in folding energy predicted by Rosetta and the DMS fitness scores for all variants.

To examine the distribution of fitness effects (DFE) of VIM-2 variants, we classified the 5291 nonsynonymous variants as having negative (<-0.7), neutral (−0.7 to 0.7) or positive (>0.7) fitness by performing Z-tests of each variant’s fitness scores against the fitness distribution of 244 synonymous variants (the null model distribution), adjusting for 5% false discovery rate using the Benjamini-Hochberg procedure. The DFE of VIM-2 variants is similar across all selection antibiotics at the highest screening concentration (128 µg/mL AMP, 4.0 µg/mL CTX, 0.31 µg/mL MEM), with ∼65% of variants being negative, ∼30% being neutral and ∼5% being positive (**Supplementary Fig. 5**). The overall DFE also agrees with observations found in DMS of other enzymes, such as *E. coli* amidase, TEM-1 β-lactamase and levoglucosan kinase^32, 36, 38, 43^.

Next, in order to determine biophysical properties that explain the fitness scores, a linear model was constructed using the fitness score of variants as the response factor and various parameters such as ASA, ΔΔ*G* predicted by Rosetta, change in amino acid volume and polarity, and the wt and variant amino acid states as predictors (**Table 1, Supplementary Tables 2** and **3** for CTX and MEM models, **Supplementary Data 3** for data used in models). We first modeled each predictor alone with the response, then selected predictors that account for at least 10% of variance in the response (R^2^>0.10) and modeled them in combinations of 2 or more as individual terms and as interacting terms. The final model accounted for the greatest amount of variance using the fewest predictors—i.e. optimized for adjusted R^2^—and combined four predictors (ASA, ΔΔ*G*, wt and variant amino acid) capable of explaining 55% of the variation in fitness scores (adjusted R^2^ = 0.55). The ASA alone can explain 21% of fitness score variation (**Table 1**) and ASA of the wt amino acid alone show a significant correlation (R^2^=0.50) to the average fitness scores of the position, with mutations at exposed residues having less deleterious fitness effects on average (**Fig. 4b**). The ΔΔ*G* explains an additional 18% (**Table 1**) and there is overall correlation between ΔΔ*G* and fitness score while individual predictions are relatively scattered, similar to previous findings that compared fitness to Rosetta folding energies or solubility scores (**Fig. 4c**)^36, 38^. Knowing the wt and variant amino acid can further explain another 10% and 5% of variation respectively^41, 51^. Thus, the results indicate structure and biophysical factors can explain the majority of fitness score tendencies.

**Table 1.**
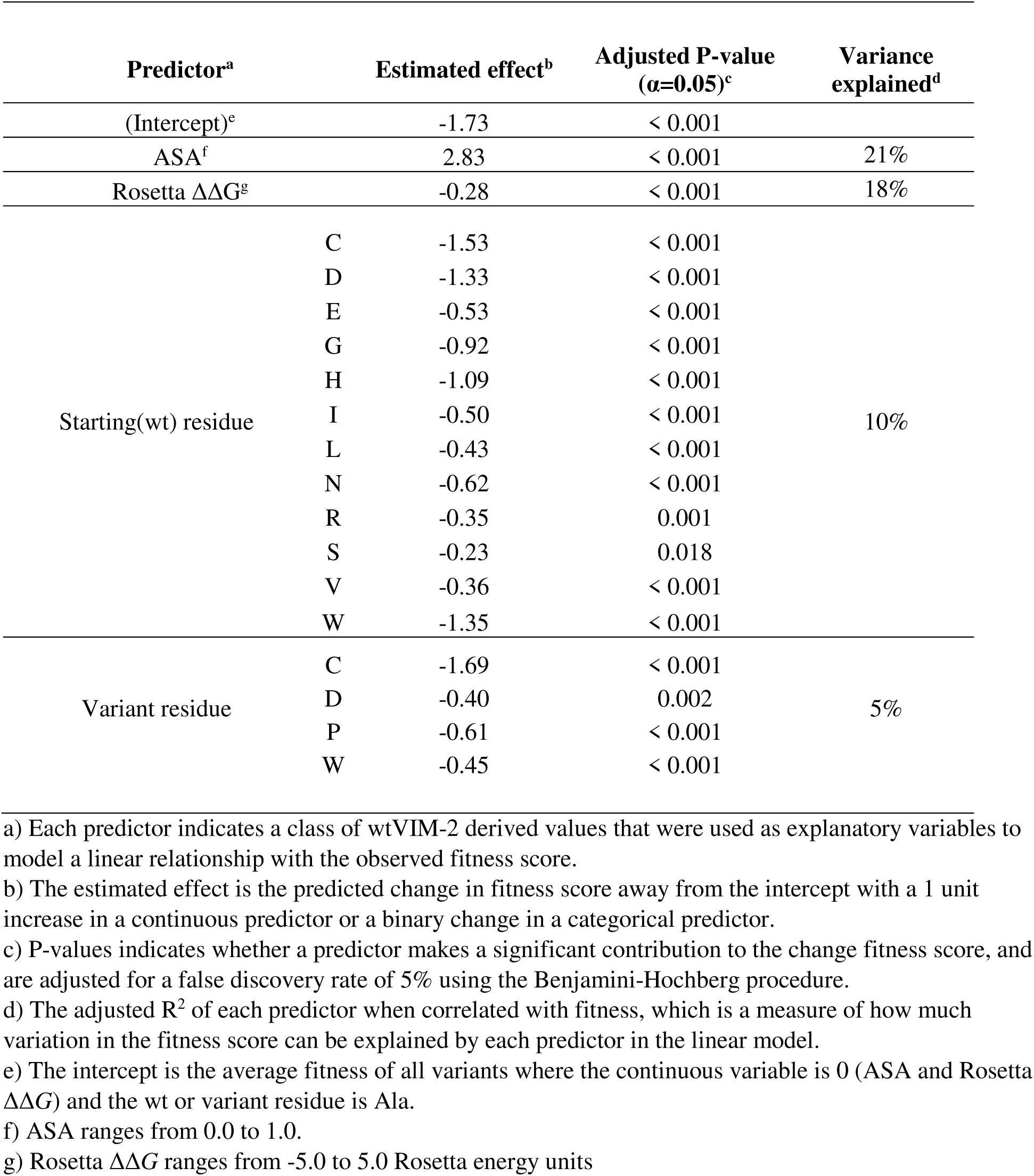
Linear model output for DMS fitness scores under 128 µg/mL AMP selection.

### Codon and amino acid optimization in the signal peptide

The first 26 residues of VIM-2 has been identified as the signal peptide^14, 15, 52^, which is a sequence used to translocate the enzyme to the periplasm, then cleaved after transport^53–56^. Our DMS data supports the known length of the signal peptide as mutations to Cys are much less deleterious before residue 26, suggesting these positions are not incorporated into the mature enzyme in the periplasm. In general, the signal peptide sequence has an amino terminal (N) region (residues 1-7) with 1 or more positive residues, a hydrophobic (H) region (residues 8-21) and a carboxy terminal (C) region (residues 22-26) that precedes the cleavage site containing a PXAXS motif (**Fig. 5a**)^53–56^. The signal peptide is conserved at 17 of 25 positions across all VIM variants, and the remaining are binary differences between the conserved sequences of the VIM-1 and VIM-2 clades (**Fig. 5a**, clades are defined by **Supplementary Fig. 6**).

**Figure 5.**
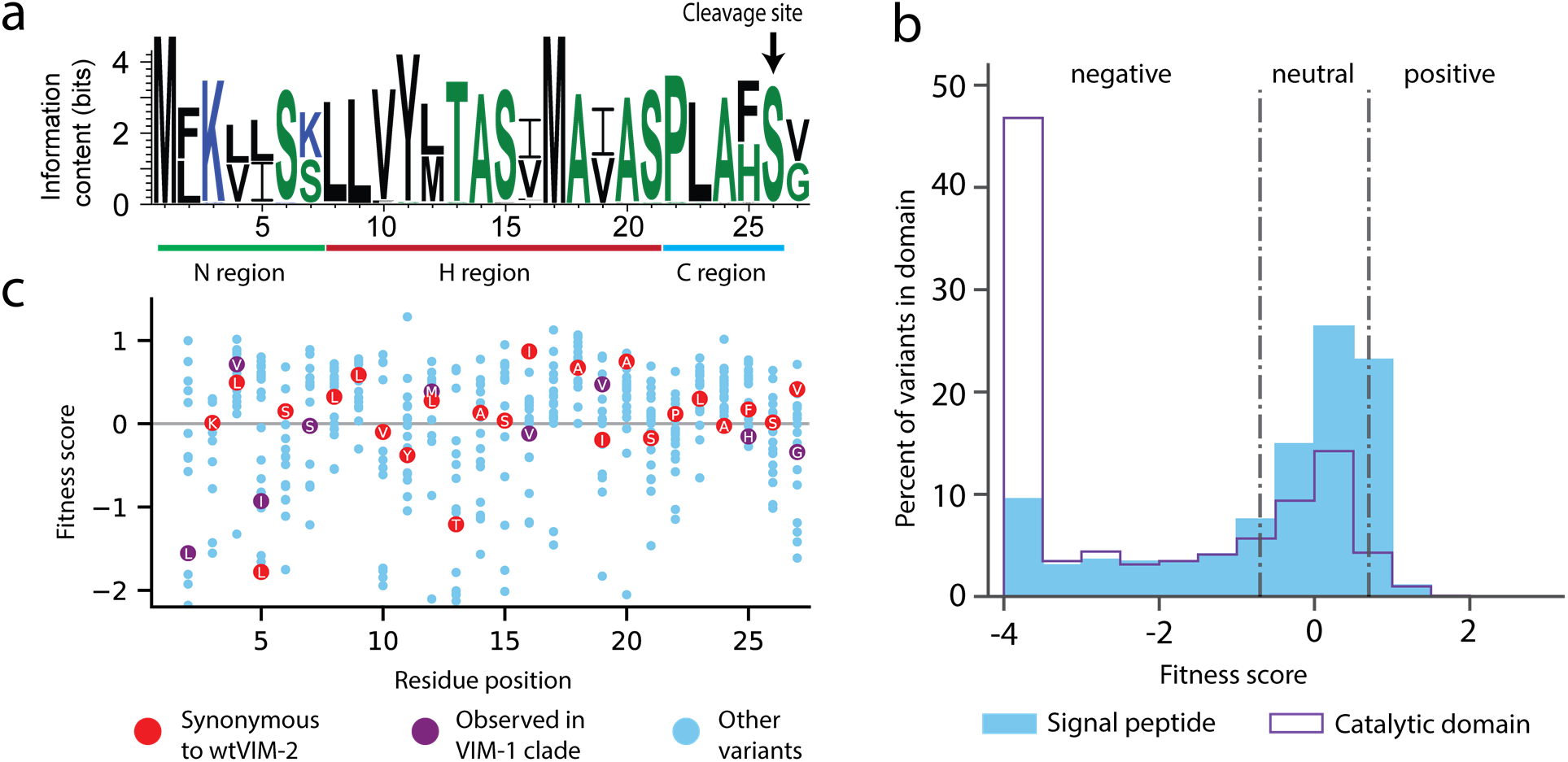
Conservation patterns and fitness scores in the signal peptide. (**a**) Sequence logo of the signal peptide region aligned across all VIM natural variants generated using WebLogo (https://weblogo.berkeley.edu/). Positions with two major naturally occurring residues are conserved differences between the VIM-1 and VIM-2 clades (clades are defined in **Supplementary Fig. 6**). (**b**) The distribution of fitness effects of all DMS variants, separated into signal peptide variants and catalytic domain variants. (**c**) DMS fitness scores of all variants at each position of the signal peptide. Synonymous variants of wtVIM-2 and conserved variants observed in the VIM-1 clade are highlighted as labelled circles.

Mutations in the signal peptide are generally tolerated (64% of nonsynonymous mutations are neutral with 128 µg/mL of AMP) and even beneficial (9.5% of mutations are positive, **Fig. 5b**), which is consistent with a previous DMS study with TEM-1^36^. In the N-terminal region, mutations to Lys3 are especially deleterious, likely due to the importance of a net positive charge in the N-region for efficient translocation^53, 57, 58^. In contrast, Lys7 is tolerant to substitution—in fact, half of the natural VIM variants have a Ser at this position—suggesting that Lys3, rather than Lys7, is critical for translocation. In the H-region, residues Val10 through Ile16 are the most sensitive to mutation, especially when changed to a charged amino acid^36, 53^. The C-region is only strongly affected by the mutation of Leu23 to Cys or Trp, while variants at other positions are neutral, including the PXAXS motif.

Interestingly, variants with evolutionarily conserved residues in the signal peptide often do not have the highest fitness, and we note both residue level and codon level dependencies in fitness (**Fig. 5c**). At most positions, a number of variants with nonsynonymous mutations produce higher fitness than wtVIM-2. Furthermore, at some positions, variants with synonymous mutations have significantly higher (Ile16, Ala18, and Ala20) or lower fitness (Lue5 and Thr13) (**Supplementary Fig. 7b**, **Supplementary Data 4**). Overall, it appears the signal peptide is less than optimal. In terms of codon level fitness among synonymous variants, we find a significant association between a mutated codon’s change in mRNA folding energy and fitness within mutants close to the start codon, supporting previous findings that avoidance of secondary structure near the start codon is favored (**Supplementary Fig. 7c**, **Supplementary Data 5**, see methods for ‘RNA folding energy calculation’)^59, 60^. We also found that 73 of 143 ‘codon dependent variants’—where any pair of synonymous codon scores differ by more than 2.0—are within the signal peptide, further supporting the importance of codon choice near the start of the coding region^61^. The less than optimal residue level fitness of wtVIM-2 may be because we employ *E. coli* as a host while natural VIM variants are often found in *Pseudomonas*^62^, and/or the signal peptide is not selected to produce maximum expression in natural environments. It has been shown that different signal peptides produce variable expression levels and translocation rates for a given protein, both of which affect the final resistance, especially in different host organisms^23, 57, 58, 63–65^. Furthermore, the signal peptide is frequently mutated in naturally occurring VIM-type variants (see section on ‘Natural VIM variation’ below), suggesting changes in the signal peptide sequences may have played significant roles in dissemination of MBL genes to different hosts and adaptation to higher antibiotic concentrations.

### Elucidation of the role of residues in the catalytic domain

We sought to further examine the functional and structural roles of residues in the catalytic domain (positions 27-266) of wtVIM-2. We compare fitness scores between selection in 128 µg/mL and 16 µg/mL AMP, as fitness at different AMP concentrations reflect a residue’s degree of involvement in the enzyme’s stability, expression and/or catalytic activity. Selection was also performed at 25°C in addition to 37°C to examine temperature dependent mutational effects, highlighting residues involved in protein folding and stability; in general, lower temperatures are permissive to variants with poor folding and high aggregation propensity while having a uniform effect on catalytic rate. To assess the role of each residue, we classified all positions in the catalytic domain into four types: *i*) ‘tolerant’, if 75% of variants are neutral even in the most stringent condition with 128 µg/mL AMP at 37°C, *ii*) ‘essential’, if 75% of variants are highly deleterious even in the least stringent condition with 16 µg/mL AMP at 25^°^C, *iii*) ‘temperature dependent’, if the difference in the fitness score between 25°C and 37°C is more than 2.0 in either 128 or 16 µg/mL AMP, and *iv*) ‘residue dependent’, if variants are temperature independent (similar scores at the two temperature) and exhibit a range of fitness rather than being mostly neutral or negative (**Fig. 6a-b**, **Supplementary Data 6** for all classifications).

**Figure 6.**
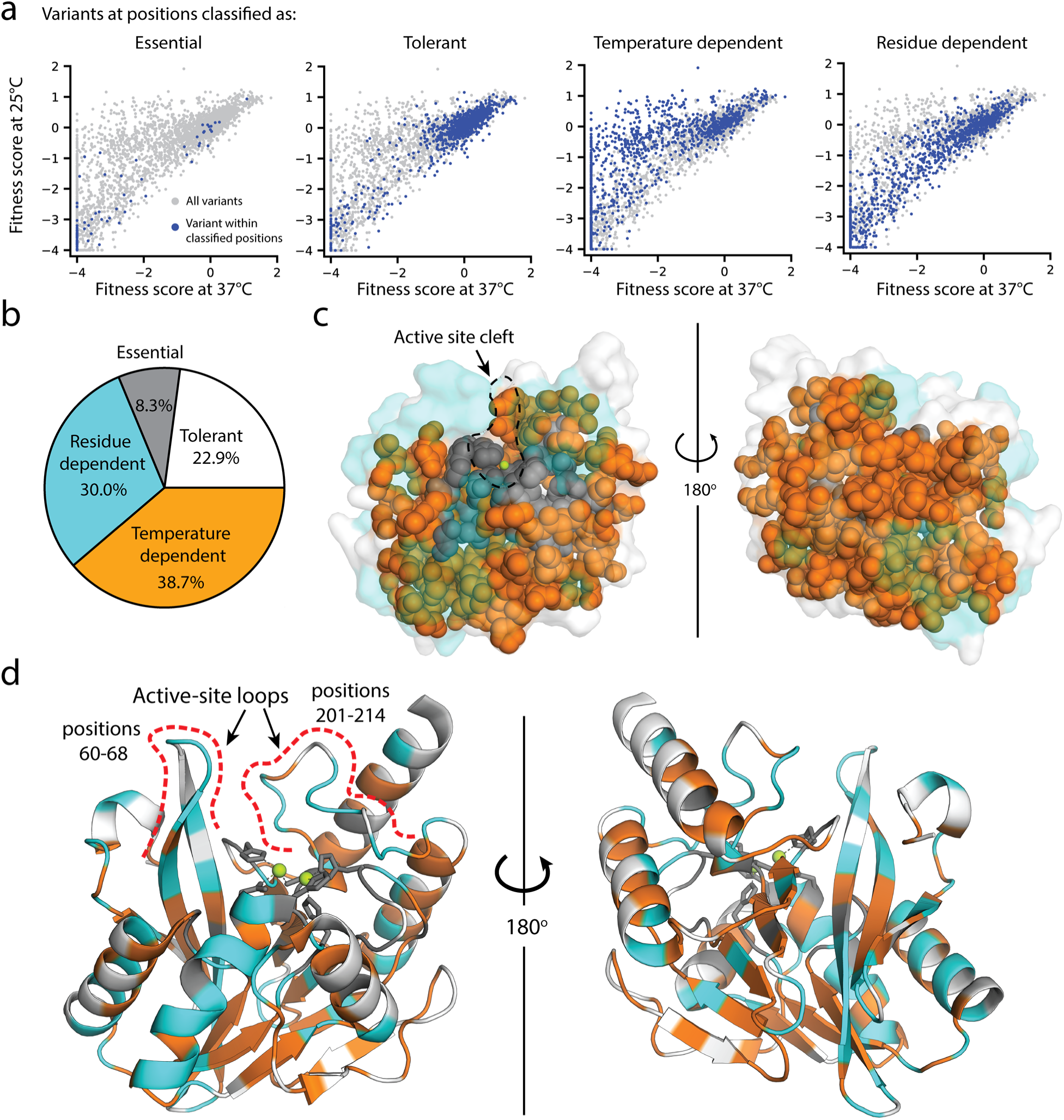
Distribution of mutational tolerance and temperature dependence of wtVIM-2 residues. (**a**) Scatterplots comparing fitness scores under selection at 25°C and 37°C (128 µg/mL AMP). Variants within the classified positions are highlighted in dark blue while all variants are plotted in grey for reference. (**b**) Proportion of residues in the wtVIM-2 catalytic domain that have been classified into each behavioral category. (**c-d**) The wtVIM-2 crystal structure (PDB: 5yd7) is colored by the behavioral classifications, the active-site zinc ions are colored in green. (**c**) View of the inner core of wtVIM-2, with essential and temperature dependent residues depicted as spheres. (**d**) Cartoon representation of wtVIM-2 with metal-binding residues shown as sticks.

As expected, the 55 ‘tolerant’ positions are scattered around the surface of the protein and are mostly solvent exposed (80% of positions have greater than 30% ASA) (**Fig. 6d**). The 20 ‘essential’ residues includes all six metal binding residues and deeply buried residues (70% have less than 5% ASA), which form the central core of the protein (**Fig. 6c**). This core is further expanded into a larger scaffold by the 93 ‘temperature dependent’ positions that are mostly buried in the structure (76% have less than 30% ASA) and largely hydrophobic—75% of the temperature dependent residues are non-polar (A, G, I, L, P, V) or aromatic residues (F, W, Y) compared to 58% for the entire catalytic domain. The 72 ‘residue dependent’ positions tend to be near the surface or at packing interfaces between α-helices and β-sheets (**Fig. 6d**).

The fitness of variants at temperature and residue dependent positions show equally strong association with the Rosetta predicted ΔΔ*G*, indicating both classes of residues have contributions to structural packing (**Supplementary Fig. 8a**). However, the location and hydrogen bonding behavior of each class of residues suggest different functional roles. Essential and temperature dependent residues display an enrichment of sidechain-backbone h-bonding relative to the proportion of h-bonding residues in each class (**Supplementary Fig. 8b**, **Supplementary Data 7**), suggesting—when combined with the formation of a hydrophobic core—these residues are largely involved in protein folding and stability^39, 46, 66^. In contrast, ‘residue dependent’ positions are prominent in the two major active site loops (14/23 positions from 60-68 and 201-214) and on packing interfaces on these loops’ distal faces from the active site. The loop holding metal-binding residues His114, His116 and Asp118, and the helix positioning the loop into the active-site (positions 112-129) are also enriched in residue dependent positions (10/15 non-metal-binding positions), suggesting possible effects on metal and substrate positioning. Thus, many of the ‘residue dependent’ positions are likely to be involved in catalysis through direct or indirect substrate interactions, and also affect the overall shape of the active site.

### Distinct recognition for different classes of β-lactam substrates

VIM-2 is known for its broad spectrum activity against all classes of β-lactam antibiotics except monobactams, but how residues achieve substrate recognition remains unknown. We examined mutations that alter substrate specificity to identify wt residues responsible for substrate recognition by comparing fitness scores between the three antibiotics (128 µg/mL AMP, 4.0 µg/mL CTX, 0.031 µg/mL MEM). First, we identified 29 ‘globally adaptive’ variants across 10 positions in the catalytic domain that increase resistance (fitness score >1) to all antibiotics (**Supplementary Data 8**). Residues at positions 47, 55, 66, 68 and 205 each give rise to at least three globally adaptive variants (24 total) while 57, 65, 115, 180 and 201 each give rise to one; 9/10 positions with globally adaptive variants are near the active site, having at least one atom within 15 Å of the active site zinc ions (**Fig. 7a**). Next, we compare fitness scores of different antibiotics in pairs, and identified variants with a change in fitness effect classifications (negative, neutral or positive) combined with a 2.0 fitness score difference between antibiotics. We identified 78 specificity altering variants across 25 positions, with 23/25 positions near the active site (**Fig. 7b**, **Supplementary Table 4** and **Supplementary Data 9** for individual specificity variants). We confirm the specificity by comparing the fitness scores with the log_2_(*EC_50 var /_ EC_50 wt_*) of the variant in the three antibiotics (**Supplementary Fig. 9**). Of the 25 positions, five are shared by both specificity and globally adaptive variants, and specificity changes are enhanced by the positive fitness at three of these positions (68, 201 and 205). However, most changes in specificity are due to decreases of fitness in one or two substrates, and only 3 variants (R205H/I/V) maintain neutral or higher fitness in all antibiotics^42, 43^. When examining the roles of these positions, we find nine ‘residue dependent’ and one ‘tolerant’ position, as expected for positions that interact with substrate rather than folding^67^. However, the other 15 positions are ‘temperature dependent’ positions, suggesting residues that are involved in substrate specificity are also embedded in the protein core.

**Figure 7.**
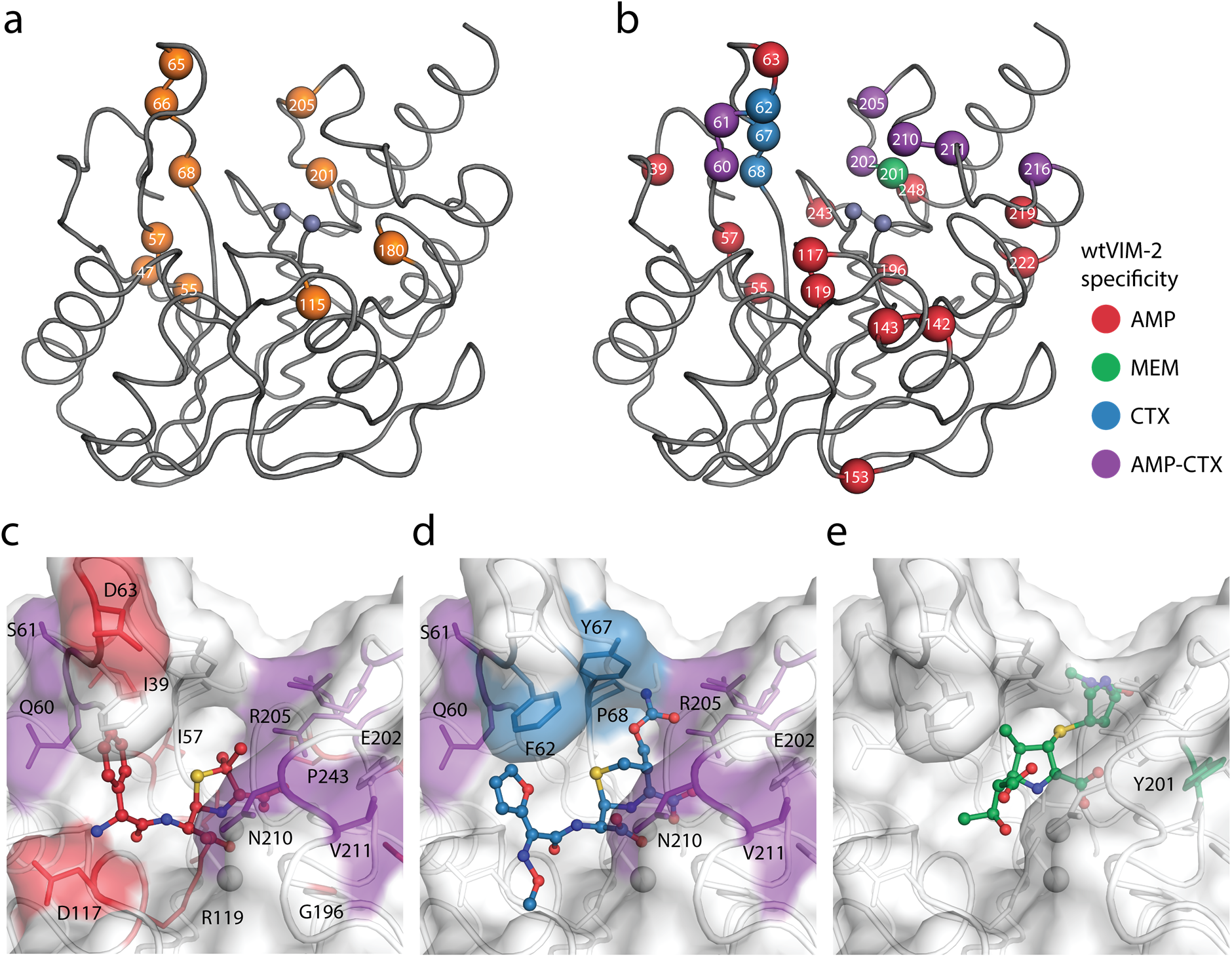
Visualization of VIM-2 specificity determining positions. The wtVIM-2 crystal structure (PDB: 5yd7) is featured in all panels, with the active site Zn ions depicted as grey spheres. (**a**) Positions with at least one globally adaptive mutation are highlighted as orange spheres. (**b**) Positions classified as being responsible for specificity towards certain antibiotics in wtVIM-2 are color coded by antibiotic and highlighted as spheres. (**c-e**). Close-up views of the specificity residues in the active site with (**c**) hydrolyzed ampicillin (PDB: 4hl2), (**d**) cefuroxime (PDB: 4rl0) and (**e**) meropenem (PDB: 5n5i) from VIM-1 and NDM-1 structures that have been aligned to the wtVIM-2 structure using the active site residues. Residues are colored by the inferred substrate specificity as in **b**). Substrates are shown in stick and ball representation.

Interestingly, there is a strong bias in specificity changes depending on mutations and their positions in the active site. In 21 of 25 positions, specificity variants decrease AMP fitness, while in 10 positions variants decrease CTX fitness and variants decrease MEM fitness at only one position (**Fig. 7b**). This bias is maintained at the level of individual specificity variants, where 96% decrease the fitness in either AMP and/or CTX—29 only decrease AMP, 33 only decrease CTX and 10 decrease both—while only 3 variants decrease MEM. The residues specific to AMP— where mutations to the residue decrease AMP fitness, but are neutral for CTX and/or MEM—are spread around the active site, including the two active site loops (60-68 and 201-214) as well as residues in the protein scaffold. In contrast, the residues specific for CTX are restricted to the two active site loops^12, 13, 68–71^; mutations in positions 62, 67 and 68 are fully specific to CTX, while positions 202, 205, 210 and 211 all have mixed specificity for CTX and AMP. To visualize residue-substrate interactions, we overlaid AMP, cefuroxime and MEM substrates in the active site of VIM-2 through alignment with VIM-1 and NDM-1 structures crystallized in complex with these substrates. Some substrate interactions are apparent from proximity, such as the packing of hydrophobic residues in loop 60-68 to the non-polar, aromatic substituents on the AMP (C6) and cefuroxime (C7) that is missing in most carbapenems (**Fig. 7c-d**). However, Glu202 and Arg205 seem to be in better position to interact with the C2 substituent of MEM and are further from AMP or cefuroxime, yet both residues are specific for AMP and CTX. Moreover, many residues in the protein scaffold that are affecting AMP specificity do not directly interact with the substrates. We hypothesize that these distant residues may contribute to solvent related phenomenon—such as displacement of solvent and/or bridging solvent with ligand to affect substrate binding energy^72, 73^—or alter protein dynamics to affect substrate specificity^22, 74–77^.

Although wtVIM-2 degrades all three β-lactams, our observations suggest that the enzyme interacts with each substrate in a different manner. AMP interacts with many residues around the active site and is the most sensitive to mutations, while CTX specificity relies exclusively on interactions with residues in the active site loops. Interestingly, MEM seems to rely on contacts shared with other antibiotics, which suggest that carbapenem resistance of VIM variants may have coevolved with other antibiotics.

### Natural VIM variation favors neutral, adaptive and specificity mutations

Currently, 56 unique VIM-type MBL sequences (including wtVIM-2) have been found on plasmids in β-lactam resistant clinical isolates (**Supplementary Table 5**)^13, 62^. The DMS data of VIM-2 enable us to characterize these naturally occurring mutations comprehensively. We classified these sequences into four clades, represented by VIM-1 (between 25-29 mutations from VIM-2 each, 45 unique mutations total), VIM-2 (1-6 mutations, 31 total), VIM-7 (70 mutations), and VIM-13 (32-33 mutations, 34 total) (**Fig. 8a**, **Supplementary Fig. 6**), with 131 unique point mutations relative to VIM-2 across 99 positions (**Supplementary Fig. 10**, **Supplementary Data 10**).

**Figure 8.**
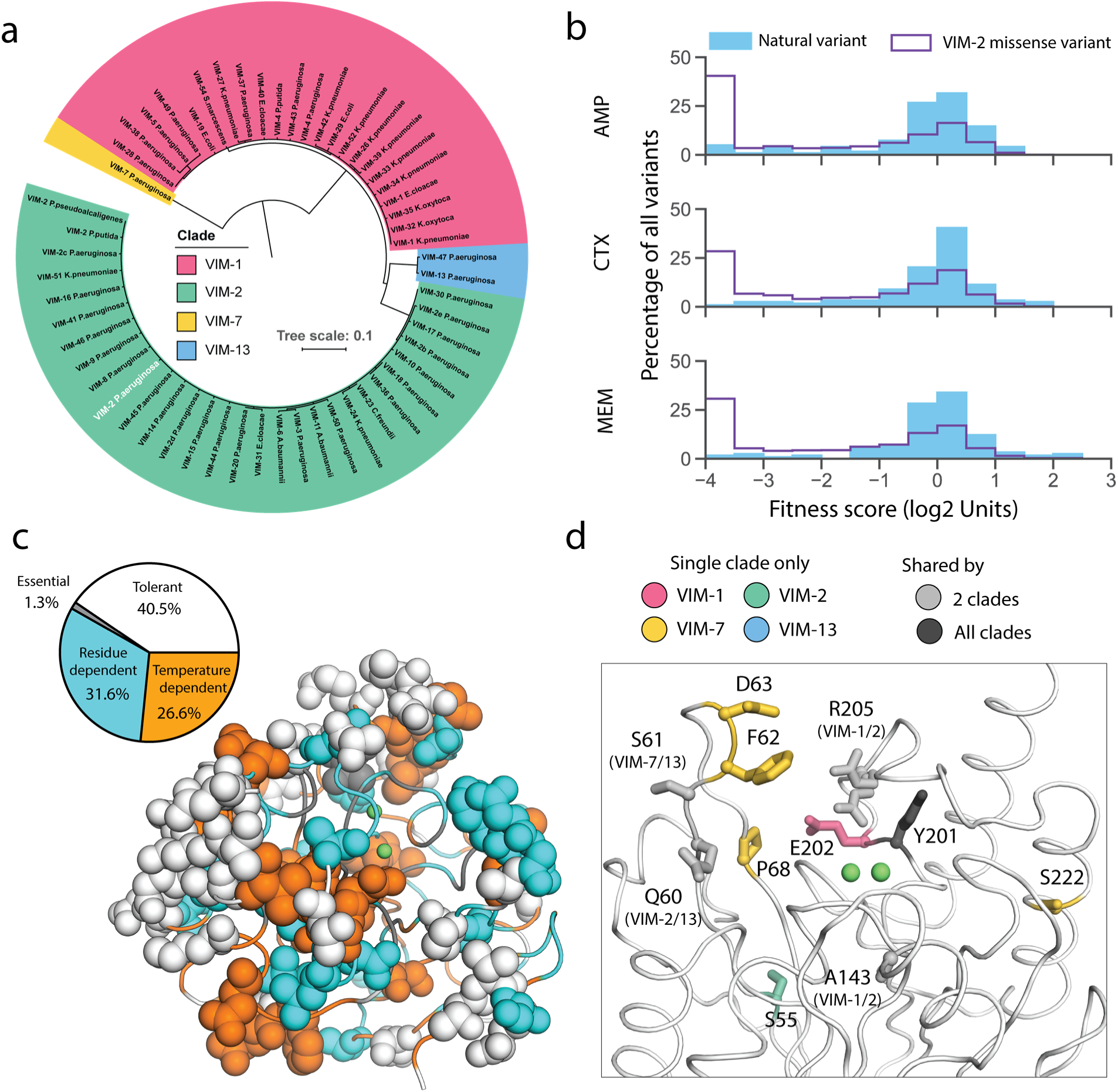
Behavior of natural VIM variants inferred from DMS fitness. (**a**) Maximum likelihood phylogenetic tree of all natural VIM variants examined in this study, colored by major clades (a larger version of the tree is presented in **Supplementary Fig. 6**). (**b**) Distribution of fitness for all unique individual mutations found in VIM natural variants compared to all missense variants measured in DMS for all 3 antibiotics. (**c**) All residues mutated in the natural variants are shown in sphere representation and colored by mutational tolerance and temperature dependence. The pie chart on the right shows the proportion of natural variant positions in each classification. (**d**) wtVIM-2 residues that are both mutated in at least one natural variant and affect specificity are highlighted as sticks, colored by the clade(s) in which the residue is mutated.

As expected, the fitness distribution of naturally occurring mutations in the VIM-type MBL for all antibiotics (128 µg/mL AMP, 4.0 µg/mL CTX, 0.031 µg/mL MEM) shows enrichment for neutral and positive mutations (**Fig. 8b**). The trend suggests that natural variants have adapted to higher resistance as 31% of all ‘globally adaptive’ catalytic domain variants in the DMS are also naturally occurring mutations, and 17% of all natural mutations have positive fitness effects for at least one antibiotic. At least 66% of variants have neutral fitness effects within each antibiotic. Natural mutations are disfavored in residues crucial for activity or stability (**Fig. 8c**): Of the 99 mutated positions, only a small proportion of ‘essential’ (1/20 positions) and ‘temperature dependent’ (21/93) positions have been mutated while large proportions of the signal peptide (20/26) and ‘tolerant’ positions (32/55) have been mutated. Interestingly, 44% (11/25) of specificity altering positions have been mutated, which suggests that VIM variants may have changed their substrate specificity during evolution (**Fig. 8d**).

However, 10% of mutations are still highly deleterious (fitness score < −2.0) in 128 µg/mL AMP (6.9% for 4.0 µg/mL CTX, 6.1% for 0.031 µg/mL MEM), indicating other factors that affect natural variation (**Fig. 8b**). These 13 deleterious mutations are spread over 11 positions, where one position is in the signal peptide, and 10 are in the catalytic domain. The signal peptide mutation (K3Q) occurs only in VIM-7 and eliminates the Lys3 critical to translocation, but this is likely neutral as VIM-7 has a S6R mutation that replaces the positive charge. Two mutations are only deleterious to AMP (Q60H, A143T), thus altering substrate specificity. Furthermore, we suspect the neutral I185V mutation (all 55 natural variants have Val185 while our wtVIM-2 has Ile185) acts as a global suppressor^78, 79^, and permit the accumulation of four mutations that are deleterious in all antibiotics (T139A, T139I, V236G and V255A) as I185V is the only other mutation in these natural variants. The remaining 6 deleterious mutations are likely neutralized by specific intramolecular epistasis, or the background dependence of mutational effects^80–84^, as these mutations occur in natural variants with at least 25 mutations relative to VIM-2. While such epistasis will hamper our ability to perfectly predict the effect of mutations, the VIM-2 dataset presented in this study contributes to further our understanding of MBL evolution, and can help orient our predictions concerning the emergence of future resistance.

## Conclusion

In this work, we report the first comprehensive mutational analysis of an enzyme in the MBL, class B β-lactamase family, one of the most important enzyme family underlying the dissemination of multi-drug resistance to pathogens. We uncover a lack of optimization in the signal peptide of VIM-2, that may be due to codon dependent RNA folding or incompatibility with the host translocation system. Such findings highlight the importance of genome and host context in resistance gene compatibility^23, 60, 65^. By performing DMS at various conditions, three different antibiotics, and two temperatures, we enhance the understanding of sequence-structure-function relationships by unveiling a set of mutations for protein stability, catalysis and substrate specificity of VIM-2. We find VIM-2’s substrate specificity altering residues to be enriched near the active site, which enables us to elucidate the molecular basis of enzyme-substrate interactions. This finding is in contrast to a previous DMS study that tested multiple substrates: specificity-altering mutations of *E. coli* amidase (amiE) were distributed across the entire structure in a global mode of specificity determination^43^. Thus, it is likely that each enzyme takes different mechanisms for recognizing diverse substrates. The monomeric MBLs have large, solvent-exposed active site clefts to recognize a wide range of substrates, while amiE which has a small, occluded active site and a hexameric quaternary structure that potentially favors controlling specificity through packing and subunit interactions^43^. However, determinants of reaction enantioselectivity in 4-OT—another enzyme active as a hexamer—are concentrated in the active site, which further highlights unique behaviors in different enzymes^85^. Regardless, understanding distinct mode of enzyme-substrate interactions will lead to design and development of new antibiotics and inhibitors to re-sensitize these enzymes. Finally, we demonstrated that VIM-type variants have been continuously evolving by enhancing their resistance as well as altering their substrate specificity in nature. The study of natural variation also reinforces the observation that mutations found to be neutral or beneficial in an experimental setting tend to be enriched in nature as well ^34, 66^. It is likely that many new VIM variants will emerge in the future, and our results will provide a valuable basis to predict likely mutations and estimate the resistance of newly found variants.

## Methods

### Materials

LB Broth, Miller (BP1426), ampicillin sodium salt (BP1760) and cefotaxime sodium salt (BP29511) were purchased from Fisher Scientific. Meropenem trihydrate (M2574) was purchased from Sigma-Aldrich (Millipore sigma). E. cloni® 10G electrocompetent cells (60061) and E. cloni® 10G chemically competent cells (60107) were purchased from Lucigen Corp. The KAPA HiFi PCR Kit (KK2102) was purchased from KAPA Biosystems Inc., the E.Z.N.A.® Cycle Pure Kit was purchased from OMEGA Bio-tek Inc. and the QIAprep Spin Miniprep Kit was purchased from Qiagen. The NextSeq 500/550 High Output Kit (300 cycles) (20024908) was purchased from Illumina Inc.

### Generation of a VIM-2 mutagenized library with all possible single amino acid substitutions

The wild-type (wt) VIM-2 gene including its signal peptide sequence from *Pseudomonas aeruginosa* was synthesized (Bio Basic Inc.) and subcloned into an in-house plasmid, pIDR2 (chloramphenicol resistance) (**Supplementary File 1**), under a constitutive AmpR promoter using *Nco* I and *Xho* I restriction enzymes (Fisher Scientific). The ATG codon in the *Nco* I site was used as the start codon. However, the cut site requires an extra G nucleotide to follow the start codon and an additional Gly (GGA codon) residue was inserted into the second position of the VIM-2 sequence; this extra Gly relative to wtVIM-2 will be labelled as G2 to distinguish it from position 2 in the wt sequence. The pIDR2 plasmid containing the wtVIM-2 gene will be referred to as pIDR2-wtVIM-2.

To generate all single amino acid variants, a library of codon mutants was made for each codon (267 positions) in the wtVIM-2 gene using restriction-free cloning (RFC)^47^ (**Supplementary Fig. 11**). We designed a forward primer for each codon position that contains a degenerate ‘NNN’ codon—using a script used to design primers for the PFunkel method^86^—and a single reverse primer, allowing a PCR to amplify part of the wtVIM-2 gene while incorporating the mutation. The PCR reaction to amplify part of the gene was done using a KAPA HiFi PCR Kit (Kapa Biosystems, Inc.) for 30 cycles of amplification each with denaturation at 98°C for 20 s, annealing at 62°C for 15 s and extension at 72°C for 15 s; 1 ng of pIDR2-wtVIM-2 was used as template in a 20 µL reaction, with 1 µL each of the forward and reverse primer (10 µM). The first PCR products were purified using E.Z.N.A.® Cycle Pure Kit (OMEGA Bio-tek, Inc.). Afterwards, 10µL of the first PCR product was then used as a primer to extend the entire plasmid, where the cycling conditions were identical to the first reaction except the extension time (90 s) and 1 ng of pIDR2-wtVIM-2 was freshly added as the template. Product from the second PCR was treated with *Dpn* I for one hour at 37°C to degrade the original wtVIM-2 plasmids, and then the amplified plasmids were purified and concentrated by the ethanol precipitation method. Subsequently, the purified plasmids were transformed into E. cloni® 10G chemically competent cells (Lucigen Corp.) using the supplier’s recommended heat-shock transformation protocol and plated on LB-Cm (containing 25 µg/mL chloramphenicol) agar plates. We then counted the number of colony forming units (CFU) obtained after the transformation for every mutagenesis library. Using CASTER^87^ and GLUE^88^, we conservatively estimated that at least 700 CFU after transformation is needed to achieve 100% coverage of all 64 codon variants per position. If a transformation met the required CFU, all colonies were collected and the plasmids were purified using QIAprep Spin Miniprep Kit (QIAGEN N.V.), while those that did not were re-transformed or re-cloned until the count was met.

### Antibiotic selection of the VIM-2 mutagenized library

Mutant libraries at individual codons were mixed into seven groups of 39 (33 for the last group) consecutive codons (see “Deep sequencing and quality control”). E. cloni® 10G electrocompetent cells (Lucigen Corp.) were transformed with 1 ng of the plasmid DNA from each of the seven groups using the supplier’s recommended electro-transformation protocol and grown overnight in 10 mL LB-Cm shaking at 30°C. We plated 1/1000 of the transformed culture on LB-Cm agar plates to estimate total CFU after transformation. Using CASTER and GLUE, it was estimated that 20,000 CFU are needed to fully cover 2496 codon mutants (64 codons × 39 positions) and all groups transformed had at least 100,000 CFU. The transformed libraries were suspended in LB media and preserved in 1 mL aliquots at −80°C in LB with 25% final volume glycerol.

Antibiotic selection was conducted in duplicate on two separate days. For each experiment, the 1 mL glycerol stock from each group was thawed and grown in 10 mL LB-Cm shaking at 30°C for 16 hrs, with optical density at 600 nm (OD600) of the cell culture reaching ∼1.5. The cultures were then diluted by 1000 fold into fresh LB-Cm and grown at 37°C for 1.5 hrs. After 1.5 hrs of growth (OD600 of the culture is between 0.01 and 0.02), 960 µL of each culture was directly introduced to 40 µL of the antibiotics at 25× concentration (final concentrations are 128, 16 and 2.0 µg/mL for AMP, 4.0 and 0.5 µg/mL for CTX, and 0.031 µg/mL for MEM, prepared in water) or water (no selection) into the wells of a 2.2mL deep-well 96 well plate, and grown for 6 hrs at 37°C. The cultures were also selected at the same AMP concentrations or grown without selection at 25°C. A culture of E. cloni® 10G electrocompetent cells transformed with pIDR2-wtVIM-2 was also grown for 6 hours at 37°C.

After placing the cultures under selection, antibiotics and DNA from lysed cells were removed by washing the selected cultures three times using a Biomek 3000 (Beckman Coulter Inc.) liquid handling robot. For each wash, the culture was centrifuged at 4000 RPM for 12 min, and the supernatant was removed using the Biomek. Subsequently, 800 µL of fresh LB was manually dispensed into the wells, the plate was sealed with plastic film, and the pellets were resuspended by vortexing. The resuspended cultures were centrifuged again for the next cycle of the wash. After the final wash, all cultures were propagated overnight shaking at 30°C in 1 mL of LB-Cm. Plasmid DNA was purified from the cultures using a QIAprep 96 Turbo Kit (QIAGEN N.V.).

### Determination of half maximal effective concentration (EC_50_)

We isolated 12 variants from codon libraries at positions 55, 62, 67, 68 and 11 variants from codon libraries at positions 205, 209, 210, 211 by transforming the libraries into *E. coli,* plating on agar plates, and picking single colonies. The identity of each variant was obtained by Sanger sequencing. The variants are grown into glycerol stocks in a 2.2 mL 96 well deep well plate; 2 single colonies of the VIM-2 WT and empty vector were also placed on this plate as controls.

The variants in the plate were then placed under the same liquid culture selection procedure as the DMS selection experiments (see “Antibiotic selection of the VIM-2 mutagenized library” above), up to the end of the 6 hrs of selection where the cell growth (OD600) was measured. The range of selection is 1 – 1024ug/mL for AMP, 0.0625 – 64ug/mL for CTX and 0.002 – 2 µg/mL for MEM, separated in 2 fold increments. All variants were also grown without antibiotics as a growth control. We calculate the *EC*_50_ by fitting **equation (2)** using the “curve_fit” function of the “Scipy.optimize” package.

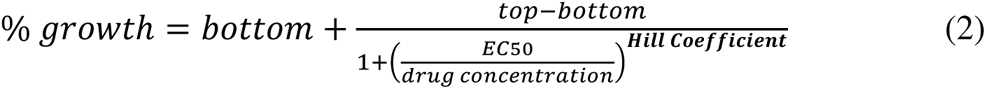

Initial estimates were 100% for top, 0% for bottom, −1.0 for the Hill coefficient and 1.0 for the *EC_50_*. In the case where a variant’s growth curve did not produce a successful fit based on the initial estimate, only the Hill coefficient and *EC_50_* were adjusted until the fit was successful. Variants where the *EC_50_* do not appear in the growth curve (stop codons, highly deleterious mutations and empty vector) could not be fitted and were excluded.

The DMS fitness scores (y-values) for each antibiotic were fitted against the *EC_50_* (x-values) using a similar sigmoidal curve in **equation (3)** with the “curve_fit” function.

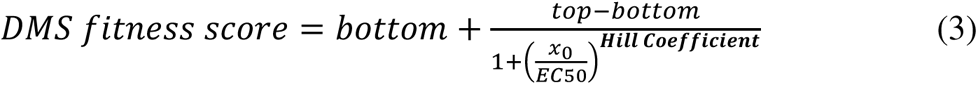

The initial guesses for top, bottom and Hill coefficient are 2.0, −4.0 and 1.0 respectively for all antibiotics. The initial estimate for x_0_, the inflection point of the curve, was 64 µg/mL for AMP, 4.0 µg/mL for CTX and 0.031 µg/mL for MEM.

The linear region of the sigmoidal curve for the DMS fitness scores was calculated using **equations (4-7)**, based on the final fitted values for each antibiotic^49^.

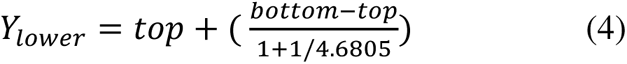

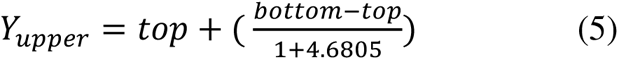

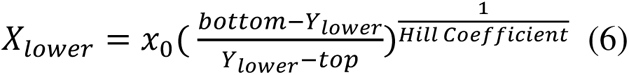

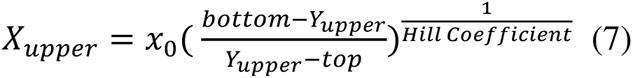

### Deep sequencing and quality control

We grouped individual codon libraries into seven groups of 39 consecutive codons (33 for the last group) so that all mutations are within a distance of 117 bp (99 bp for the last group), allowing 150 bp forward and reverse NGS sequencing reads to generate full overlap of each group. PCR amplicons of each library group and wtVIM-2 were generated using primers that flank the 117 bp region of each group, where primers have a Nextera transposase adapter sequence (Illumina, Inc.) in the 5’ overhangs. We used KAPA HiFi PCR Kit (Kapa Biosystems, Inc.) for 15 cycles of amplification each with denaturation at 98°C for 20 s, annealing at 65°C for 5 s and no extension time. The amplicons were extended by PCR again to include the sample indices (i7 and i5) and flow cell binding sequence, then sequenced using a NextSeq 550 sequencing system (Illumina, Inc.); all samples were sequenced in the same NextSeq run. The raw sequencing data can be found on the NCBI Sequencing Read Archive (SRA) (BioProject accession: PRJNA606894). Each group under each condition received between 400,000 to 1,000,000 reads, which is at least 160 reads per codon variant on average. There were 5 samples that received as low as 100,000 reads in one of the two replicates, but the correlation of fitness scores between both replicates still showed an R^2^ of at least 0.72 and the scores were retained.

To process the NGS data, including merging paired-end reads, quality filtering, variant identification and fitness score calculation, we use a set of in-house Python scripts (https://github.com/johnchen93/DMS-FastQ-processing). Paired-end reads in FastQ format were first merged using a Python script, where quality (Q) scores of matching read positions were combined using a posterior probability calculation to obtain posterior quality (Q) scores, measured on the Phred scale for sequencing quality^89^. In the case of a base mismatch between the forward and reverse reads, the base was taken from the read with the higher Q score at the position.

Reads that had more than 20 base mismatches between the forward and reverse reads, or that had any bases with a posterior Q score less than 10 were discarded. It was found that above a Q score of 10 or more, the average proportion of sequencing errors per position stabilized and no sizeable reduction of sequencing errors can be obtained by Q score cut-offs (**Supplementary Fig. 12a**). Usually, 75-85% of all reads passed these filters. Additionally, the expected number of errors per read was calculated from adding the error rates calculated from the posterior Q Scores of every position in the entire read^89^. Reads that had an expected number of error greater than 1 would also be discarded, however no reads exceeded this limit after the previous filters.

### Variant identification and noise filtering

Once forward and reverse reads were merged and filtered by read quality, codon mutants were identified and counted, then aggregated into amino acid (or stop codon) variants. Codon mutations were identified by comparing to the wtVIM-2 sequence as a reference. Since we only intended for single codon mutants in the library, any sequence with mutations in more than one codon was discarded, leading to retention of 80-90% of the filtered reads.

To exclude variants that may be due to sequencing errors alone, we estimated the expected frequency of each variant generated by sequencing errors and excluded variants that have less than 2× the expected frequency in the non-selected library. Using sequencing data from wtVIM-2, we calculated the error rates that originates from culture growth, sample preps (PCRs) and sequencing. The error rates at each position was calculated by dividing the errors observed by the total number of reads at that position (**Supplementary Fig. 13d**). The mean of the distribution of the positional errors was used as the estimate for error rates (0.072%) in all positions across the VIM-2 gene. The proportion of each type of nucleotide error (A>T, A>C, A>G, etc.) was calculated to estimate the likelihood of each type of nucleotide error given a starting nucleotide (**Supplementary Fig. 12b**).

We made the observations that 1) sequencing errors in wt sequences will generate single codon mutants, but errors in single codon variants are most likely to be turned into double codon mutants (**Supplementary Fig. 13a**) 2) ∼5-10% of reads in each library group were occupied by wt sequences while other variants are rarely higher than 0.5% (**Supplementary Fig. 1b**) and 3) single nucleotide sequencing errors are the most abundant type of errors affecting up to 10% of all reads, while higher numbers of sequencing errors are nearly negligible (**Supplementary Fig. 13c**). In summary, single nucleotide errors on wt sequences are the main source of single codon variants arising from sequencing errors. Thus, we first calculated the chance of each of the 64 codons to mutate into the 9 adjacent codons by single nucleotide sequencing errors using **equation (8)** (**Supplementary Fig. 13e**).

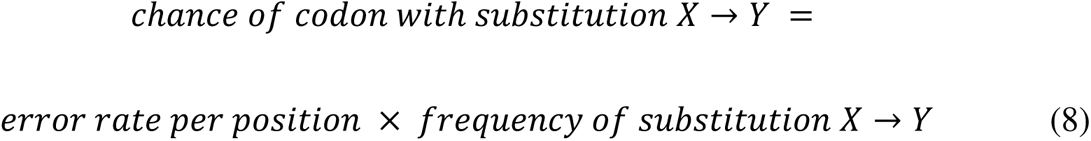

For example, to gauge how often AAA gets mutated to GAA by chance, we multiplied the per position error rate by the proportion of G mutations when starting from an A, leading to 0.0719% × 70.7% = 0.000719 × 0.707 = 0.00051. This means for each 100,000 wt reads that has an AAA codon at a given position, we expect 51 GAA mutants on average that arise by chance at the same position. We calculated expected codon error frequencies from every codon, then summed the expected error frequencies of the codons mutant for each amino acid variant (**Supplementary Fig. 13f**). The error frequency is multiplied by the count of the wt reads in each non-selected library group to arrive at an expected error count for that group. Subsequently, we compared the observed count of amino acid (or codon) variants in the non-selected library to the expected count from errors alone and we accepted a variant as truly existing if the observed count is at least twice the expected count from errors. In addition, because our filtering method only accounts for the 9 codons with a single mutation relative to the wt codon, we also applied a count cut-off of 5 for all variants to reduce noise by excluding very low count data.

### Fitness Score Calculations

The fitness score of each variant was calculated according to **equation (1)** (see main text). To calculate the fitness score of a given amino acid (or codon) variant, the read count of the variant was first normalized to frequencies within the non-selected or selected library group. Variants that exist in the non-selected library but disappear in the selected library were interpreted to have been removed by antibiotic selection, and were given a dummy count of 1 to emulate the minimum frequency observable for that variant. The frequency of the variant after selection was divided by the frequency of the same variant in the library grown without selection for an enrichment ratio; synonymous codon variants were also considered as variants rather than wt during scoring. The variant enrichment ratio was then normalized to the enrichment ratio of the wt. The final score was expressed in Log_2_ units, and scores were calculated separately across the 7 groups and separately for each replicate.

When combining all data across the 7 groups, we subtracted the mean fitness scores of all synonymous variants in each group from all variants of that group to center the mean fitness of synonymous variants at a fitness score of 0. To combine scores from replicates, we simply averaged the fitness scores across the two replicates, and take the single score if only one replicate contained the variant above noise in the non-selected library.

### Fitness effect classification

To classify each amino acid variant as positive, neutral or negative for each of the three selection antibiotics, we use the score of each variant in a two tailed z-test on a normal distribution (null-model) with the same mean and standard deviation as our synonymous distribution (244 synonymous variants total). The p-values were then FDR corrected to an α of 0.05 using the Benjamini-Hochberg procedure; only missense variants were tested and the total number of tests was 5291. The variants with scores that are significantly different from the synonymous distribution after FDR correction are then classified as ‘positive’ if their score is greater than the synonymous mean and ‘negative’ if the score is less, while the remaining variants are classified as ‘neutral’.

### Linear Model of DMS Scores with various predictors

We generated a linear model in R using a combination of terms to try and find properties that best explain the behavior we see in the DMS fitness scores. Using the fitness score as a response, we tried using **1)** wild-type (wt) amino acid **2)** variant (var) amino acid **3)** accessible surface area (ASA) of the residue calculated from the crystal structure of wt VIM-2 (PDB: 4bz3) using ASA view^90^ **4)** change in amino acid volume^91^ (Δvolume = volume_var_-volume_wt_) **5)** change in amino acid polarity (hydrophathy index^92^) (Δpolarity = Δpolarity_var_ – Δpolarity_wt_) **6)** distance of the alpha carbon of each residue in the crystal structure to the active site water held between the Zn ions **7)** Rosetta predicted stability change between the variant and the wt (ΔΔ*G* = Δ*G*_var_-Δ*G*_wt_) (also see “Rosetta ΔΔ*G* Calculation” below) and **8)** BLOSUM62 score for the substitution from wt to variant. Only variants from positions observable in the crystal structure were modelled (positions 32 to 262), and synonymous variants were excluded.

All parameters were first modelled individually as predictors with DMS fitness score as the response, and the predictors with R^2^ higher than 0.10 are then modelled in combinations of two or more until the combination with the least predictors and the highest adjusted R^2^ was found. Predictors with R2 less than 0.10 are also retried in combination with the best predictors when optimizing for adjusted R^2^. Interaction between predictors were tested, but they did not improve adjusted R^2^ and were excluded for the sake of simplicity. The relative contribution of each term to the overall adjusted R^2^ were calculated using the R package ‘RelImpo’, using the ‘lmg’ method ^93^.

The final equation of the linear model is shown in **equation (9)** where the fitness score of a given variant is the additive combination of the model intercept β_0_ and the various properties and the coefficients of the properties (e.g. β_ASA_ and ASA) plus a random error term ε.

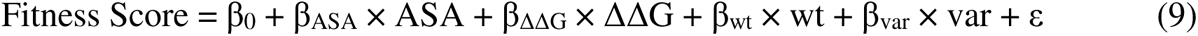

The categorical predictors wt and var are simplified in the equation and each is actually a collection of terms in the model, where every amino acid is a single binary term represented by 0 or 1 such that 1 indicates the presence of the amino acid. For example, the variant Q60V has Q = 1 for wt and V = 1 for var, while all other wt and var amino acids are set to 0. Ala is not present as one of the estimates in either wt or var, because they are used in calculating the intercept; this is the mean fitness of all data points where either wt = 1 or var = 1 for alanine.

### Rosetta ΔΔG calculation

To estimate the effects of each VIM-2 variant on the stability of the protein, we used the Rosetta “ddg_monomer” application to calculate the folding energy of a monomeric protein crystal structure. Rosetta was run on the Compute Canada server Cedar using a Rosetta 3.8 installation. Following the ‘ddg_monomer’ documentation, the VIM-2 structure (PDB: 4bz3) was first processed using ‘preminimize’ to pre-optimize the packing of the crystal structure and generate a constraints file. Then, all single amino acid variant structures and the wt structure at each position were simulated 50 times each using ‘ddg_monomer’, configured to protocol 16 as specified in Kellog *et al.* 2011^94^ while using Talaris 2014 as the scoring function. We store the simulated structures (variant and wt) as PDB files and scored them using the Rosetta ‘score’ function with Talaris 2014 weights to obtain the predicted Δ*G* in Rosetta Energy Units (REU). We average the predicted Δ*G* of all 50 replicates of each variant or wt. The ΔΔ*G* is calculated using **equation (10)** as the difference in average Δ*G* between variant and wt at the same position.

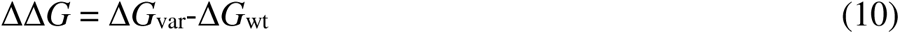

### RNA folding energy calculation for single codon mutants

The RNA folding energy contribution of the 5’ UTR and signal peptide region is calculated according to a previously described method^59^. The Δ*G* of folding of the 5’ UTR and signal peptide is calculated using **equation (11)**.

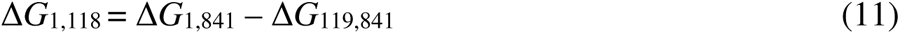

Each Δ*G* term is calculated using the NUPACK software package^95^, using the ‘pfunc’ program which calculates the Δ*G* of all RNA secondary structures from the partition function. All folding energies were calculated using default conditions, with [Na+] = 1 M and T = 37°C, and the results are in units of kcal mol^-1^. The subscripts for each Δ*G* term indicates the first and last nucleotide position in the transcript that is used for calculating the Δ*G* of folding, respectively. Thus, Δ*G*_1,841_ represents the calculated folding energy of the full transcript (including the 5’ UTR and coding region, but excluding the 3’ UTR), while Δ*G*_119,841_ is the folding energy of the transcript after the signal peptide. The interpretation of Δ*G*_1,118_ is that it is the folding energy contribution of the RNA transcript up to the end of the signal peptide, including energy from non-local interactions with downstream parts of the transcript but excluding folding energy of interactions exclusively within positions downstream of the signal peptide. The DNA sequence of the wtVIM-2 transcript used to calculate the folding energies is shown below, with positions 1-118 italicized and the translated signal peptide region underlined for clarity; the sequence is converted to RNA for calculation.

*5’-CTGATAAATGCTTCAATAATATTGAAAAAGGAAGCCC<u>ATGGGATTCAAACTTTTGAGTAAGT TATTGGTCTATTTGACCGCGTCTATCATGGCTATTGCGAGCCCGCTCGCTTTTTCC</u>*GTAGA TTCTAGCGGAGAATATCCGACAGTCAGCGAAATTCCGGTCGGGGAGGTCCGGCTTTACCAGATTGCCGATGGTGTTTGGTCGCATATCGCAACGCAGTCGTTTGATGGCGCAGTCTACCCGTCCAATGGTCTCATTGTCCGTGATGGTGATGAGTTGCTTTTGATTGATACAGCGTGGGGTGCGAAAAACACAGCGGCACTTCTCGCGGAGATTGAGAAGCAAATTGGACTTCCTGTAACGCGTGCAGTCTCCACGCACTTTCATGACGACCGCGTCGGCGGCGTTGATGTCCTTCGGGCGGCTGGGGTGGCAACGTACGCATCACCGTCGACACGCCGGCTAGCCGAGGTAGAGGGGAACGAGATTCCCACGCACTCTCTTGAAGGACTTTCATCGAGCGGGGACGCAGTGCGCTTCGGTCCAGTAGAACTCTTCTATCCTGGTGCTGCGCATTCGACCGACAACTTAATTGTGTACGTCCCGTCTGCGAGTGTGCTCTATGGTGGTTGTGCGATTTATGAGTTGTCACGCACGTCTGCGGGGAACGTGGCCGATGCCGATCTGGCTGAATGGCCCACCTCCATTGAGCGGATTCAACAACACTACCCGGAAGCACAGTTCGTCATTCCGGGGCACGGCCTGCCGGGCGGTCTTGACTTGCTCAAGCACACAACGAATGTTGTAAAAGCGCACACAAATCGCTCAGTCGTTGAGTAA-3’

**Equation (12)** is used to calculate the ΔΔ*G* of folding upon codon mutations in the signal peptide.

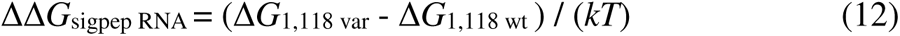

Δ*G*_1,118 wt_ is the folding energy of the wtVIM-2 signal peptide sequence shown above, while Δ*G*_1,118 var_ is the folding energy of the signal peptide sequence with a single codon mutation. The difference in folding energy is normalized to the thermal energy factor *kT*, where *k* = 0.0019872041 kcal mol^-1^ K^-1^ and *T* = 310.15 K (37°C).

### Identification of critical residues and temperature dependence

We classify the role of residues in wtVIM-2 by examining the DMS fitness scores of selection conducted at 128 µg/mL and 16 µg/mL AMP at 37°C and 25°C (4 data sets), excluding signal peptide residues 1-26. Residues are classified as ‘essential’ when >75% of variants are below a fitness score of - 2.0 when selected at the least stringent condition of 16 µg/mL and 25°C. Residues are classified as ‘tolerant’ when >75% of variants are above a fitness score of −1.0 when selected at the most stringent condition of 128 µg/mL and 37°C. Residues are classified as ‘temperature dependent’ when the fitness score of 25°C selection is higher than 37°C selection of the same AMP concentration by at least 2.0, for two or more variants at either 128 µg/mL or 16 µg/mL. The ‘temperature dependent’ classifications overwrite ‘essential’ or ‘tolerant’ classifications (4 occurrences total). Residues that do not fall into the three other classifications are defined as ‘residue dependent’.

### Detecting hydrogen bonds in the wtVIM-2 crystal structure

Potential hydrogen bonding pairs in the wtVIM-2 crystal structure (PDB: 5yd7, chain A only) were extracted using the ‘Polarpairs’ script in PyMol (https://pymolwiki.org/index.php/Polarpairs). The script filters for pairs of h-bond donor and acceptor atoms within a defined distance and h-bond angle; we set the distance limit to be within 3.6 Å and the h-bond angle to be greater than 63°. The script returns atom indices, which were converted to PBD atom IDs using pymol’s built in ‘id_atom’ function. The atoms were then extracted from the 5yd7 pdb file using the atom IDs, and the atom’s name was used to determine if the h-bond was formed between backbone atoms only (‘N’ and ‘O’ atoms indicate backbone amides and carbonyls, respectively), between sidechain atoms only or between backbone and sidechain atoms. All extracted h-bonds can be found in **Supplementary Data 7**.

### Analysis of specificity variants

We define variants with altered specificity by filtering for variants with a change in fitness effect classifications (positive, neutral and negative, defined in “Fitness effect classification”) as well as a fitness score difference of 2.0 between at least 2 of the 3 antibiotics being compared.

Specificity positions are visualized on the wtVIM-2 structure (PDB: 5yd7) using PyMol, with substrates overlaid from aligned MBL homolog structures (AMP - NDM-1(PDB:4hl2), cefuroxime – NDM-1(PDB:4rl0), MEM – VIM-1 (PDB:5n5i)). To overlay the substrates, the 6 metal binding residues (structure positions 114, 116, 118, 179, 198, 240 for VIM-1/VIM-2 and 120, 122, 124, 189, 208, 250 for NDM-1) and the 2 active-site Zn ions were selected from each structure, and the PyMol ‘align’ function was used with the VIM-2 active-site as the target object and the other structure’s active-site as the mobile object. When structures have more than one chain (PDB: 5yd7, 4hl2 and 4rl0), only chain A was used in the alignment. The protein portions of the homolog structures are hidden after alignment, to visualize just the substrates with the wtVIM-2 structure.

### Collection of naturally occurring VIM variants

Amino acid sequences of naturally observed VIM variants were extracted by performing a BLASTP search of the NCBI non-redundant protein database ^96^, using the protein sequence of our in-house VIM-2 with Gly removed from the 2nd position, identical to the VIM-2 discovered in *Pseudomonas aeruginosa* isolates (UniProt accession: A4GRB6). We retained all BLASTP results with at least 70% identity and >90% query coverage. We also retrieved protein sequences from the Comprehensive Antibiotic Resistance Database (CARD) ^62^. All sequences from both sources were merged and sequences that are exactly identical in length and sequence were combined, while sequences that are less than 250 or greater than 290 residues were excluded. Recombinant variants VIM-12 and VIM-25 were excluded from this analysis, as well as VIM-14 (UniProt accession: Q6GUL7) which is a member of the VIM-1 clade despite being labelled as both VIM-11 and VIM-14 (UniProt accession: A0SWU7) (both are in the VIM-2 clade).

To identify all mutations different between wtVIM-2 and the 55 other variants, a multiple sequence alignment (MSA) was constructed using the MUSCLE method in MEGA 7 (version 7.0.26)^97^. Equivalent positions bearing a different amino acid from VIM-2 were identified as mutations, while deletions are ignored (1 in VIM-1, 1 in VIM-7, 4 in VIM-18). The MSA was also used to generate a maximum likelihood phylogenetic tree using MEGA 7 (default settings). The tree was used to identify separate VIM clades which were labelled using the VIM variant with the lowest number in the clade (VIM-1, VIM-2, VIM-7, VIM-13).

