## Supplementary material for "Comprehensive exploration of the translocation, stability and substrate recognition requirements in VIM-2 lactamase": Supplementary Figures and Captions v3_final.pdf

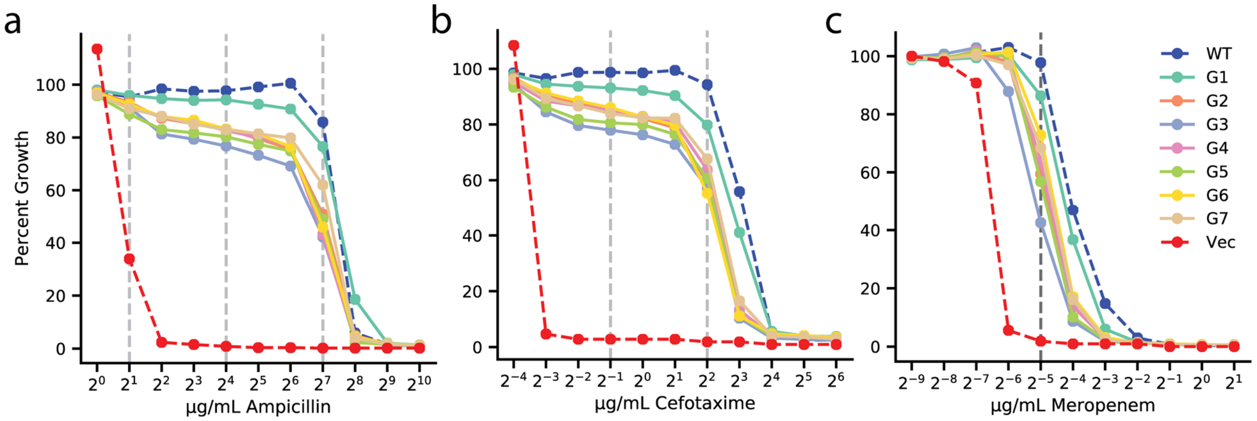


**Supplementary Figure 1. Antibiotic dose-response curves of VIM-2 libraries for all antibiotics.** (**a**) Ampicillin. (**b**) Cefotaxime. (**c**) Meropenem. Y-axis indicates percent growth of *E. coli* transformed with plasmid library groups relative to no selection. The range of antibiotics tested is 1.0-1024 µg/mL for AMP, 0.625-64 µg/mL for CTX, and 0.002-2.0 µg/mL for MEM. The dashed vertical lines indicate concentrations that were used in DMS.

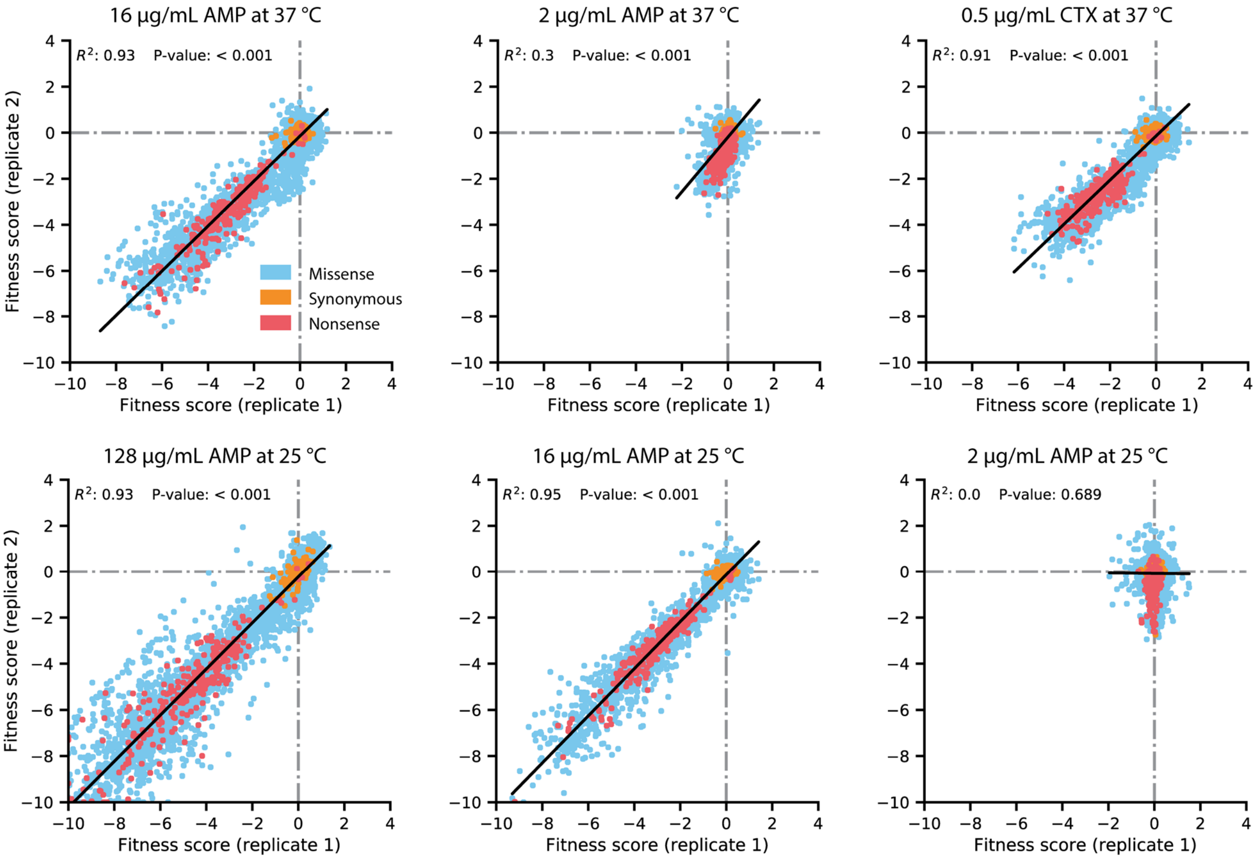


**Supplementary Figure 2. Replicate correlation of fitness scores in the DMS experiments.** The antibiotic concentration and growth temperature used for each experiment are labelled above each plot. Data points are colored by the type of amino acid mutation, as described in the legend in the top-left plot. The black line indicates the line of best fit for a linear regression between the replicates, with the $R^2$ and P-value of the regression displayed near the top of each plot.

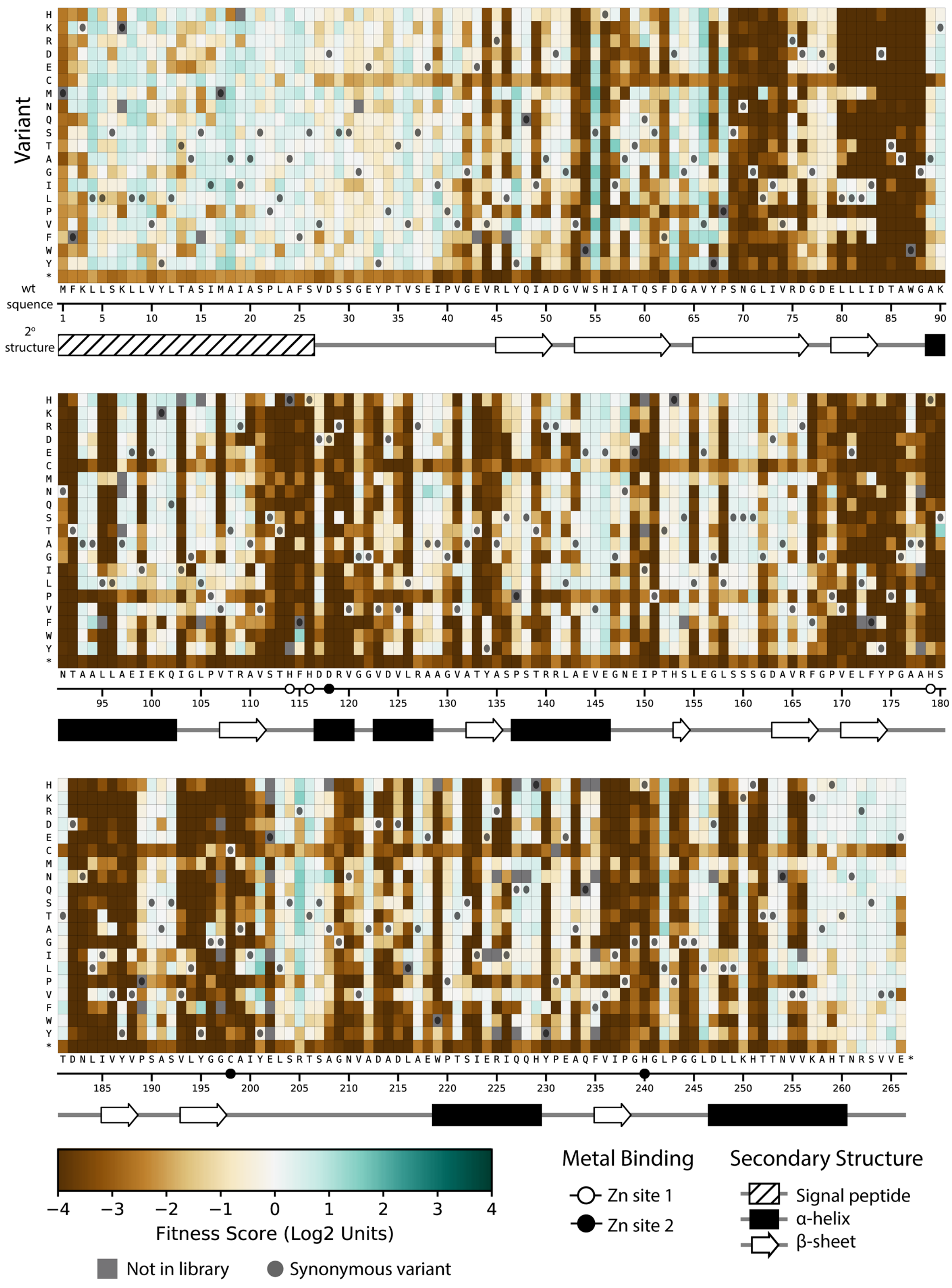
Variant
H K R D E C M N Q S T A G I L P V F W Y *
wt squence
M F K L L S K L L V Y L T A S I M A I A S P L A F S V D S S G E Y P T V S E I P V G E V R L Y Q I A D G V W S H I A T Q S F D G A V Y P S N G L I V R D G D E L L L I D T A W G A K
1 5 10 15 20 25 30 35 40 45 50 55 60 65 70 75 80 85 90
2° structure
N T A A L L A E I E K Q I G L P V T R A V S T H F H D D R V G G V D V L R A A G V A T Y A S P S T R R L A E V E G N E I P T H S L E G L S S S G D A V R F G P V E L F Y P G A A H S
95 100 105 110 115 120 125 130 135 140 145 150 155 160 165 170 175 180
T D N L I V Y V P S A S V L Y G G C A I Y E L S R T S A G N V A D A D L A E W P T S I E R I Q Q H Y P E A Q F V I P G H G L P G G L D L L K H T T N V V K A H T N R S V V E *
185 190 195 200 205 210 215 220 225 230 235 240 245 250 255 260 265
−4 −3 −2 −1 0 1 2 3 4
Fitness Score (Log2 Units)
Not in library
Synonymous variant
Metal Binding
Zn site 1
Zn site 2
Secondary Structure
Signal peptide
α-helix
β-sheet

**Supplementary Figure 3. Fitness of all VIM-2 single amino acid variants under 4.0 µg/mL CTX selection.** Each cell in the heat map represents the fitness score of a single amino acid variant. Synonymous variants are indicated as black circles and variants that are not present in the library are in grey. The x-axis under the heatmap indicates the wt residue and position (the 6 active site metal binding residues are highlighted as circles), while the y-axis indicates the variant residue at that position. The secondary structure of the wtVIM-2 crystal structure (PDB: 4bz3) is displayed below the heatmap.

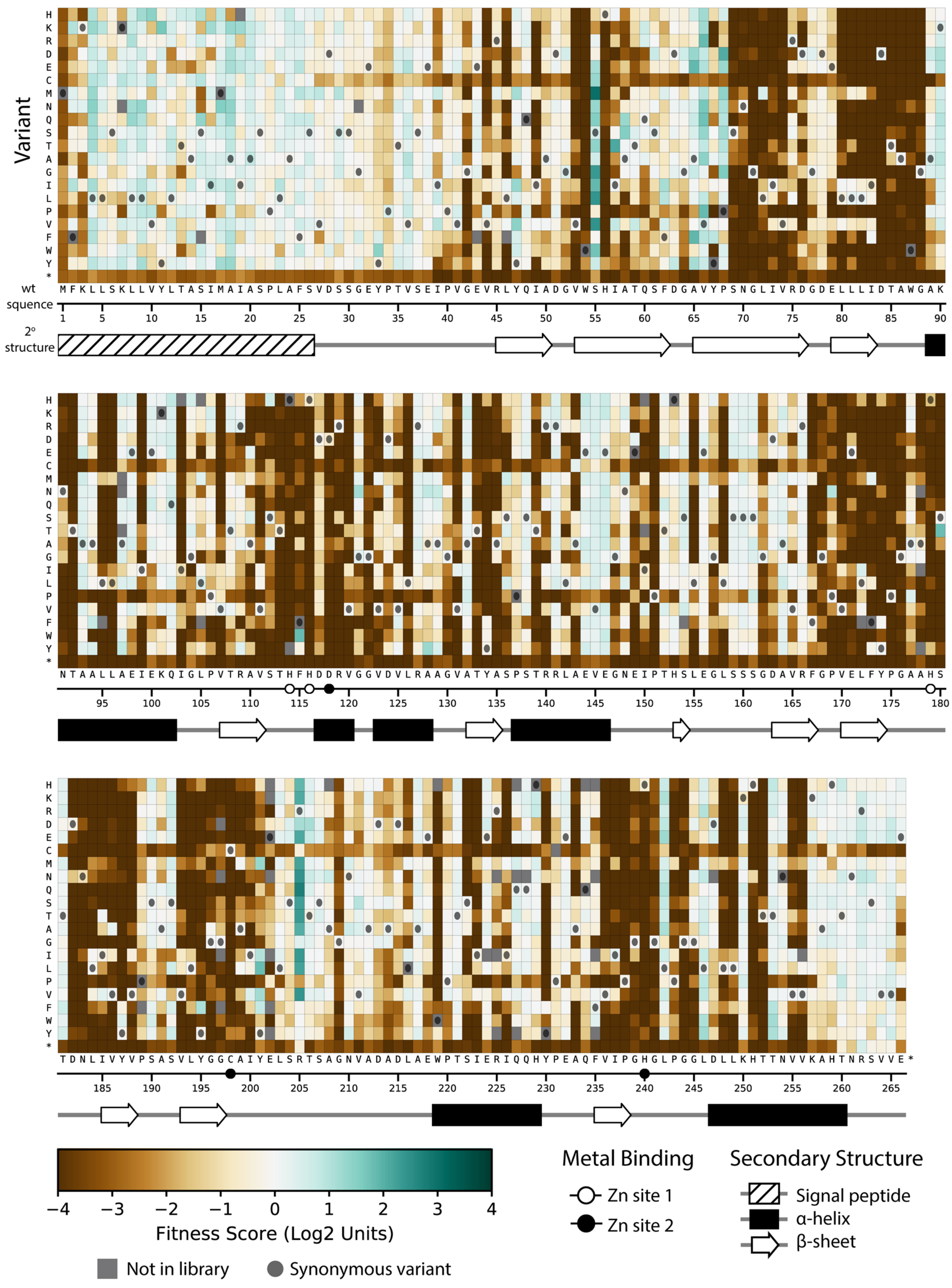
Variant
wt squence
2° structure
Fitness Score (Log2 Units)
Not in library
Synonymous variant
Metal Binding
Zn site 1
Zn site 2
Secondary Structure
Signal peptide
α-helix
β-sheet

**Supplementary Figure 4. Fitness of all VIM-2 single amino acid variants under 0.031µg/mL MEM selection.** Each cell in the heat map represents the fitness score of a single amino acid variant. Synonymous variants are indicated as black circles and variants that are not present in the library are in grey. The x-axis under the heatmap indicates the wt residue and position (the 6 active site metal binding residues are highlighted as circles), while the y-axis indicates the variant residue at that position. The secondary structure of the wtVIM-2 crystal structure (PDB: 4bz3) is displayed below the heatmap.

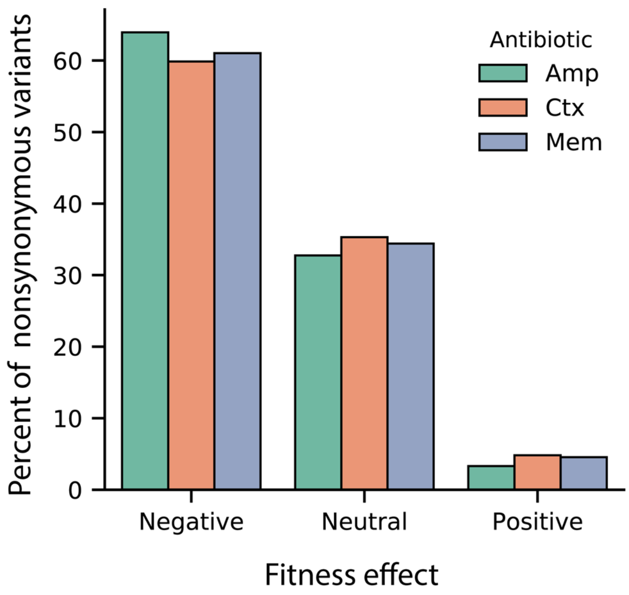


**Supplementary Figure 5. Proportion of fitness effects for VIM-2 nonsynonymous variants selected in AMP, CTX and MEM.** The proportion of variants with positive, neutral or negative fitness effects as classified for 128 µg/mL AMP, 4.0 µg/mL CTX and 0.031 µg/mL MEM**.**

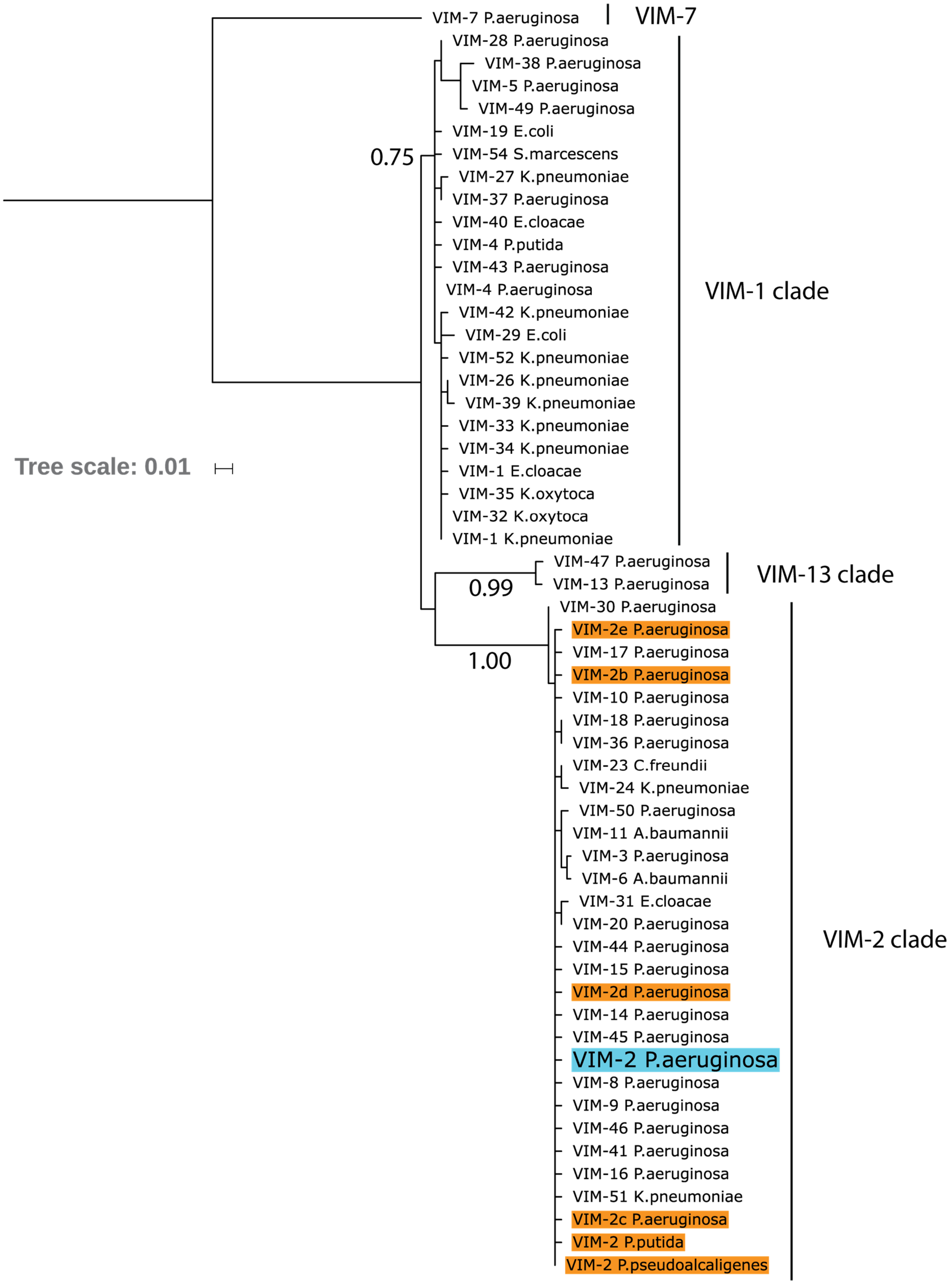

VIM-7 P.aeruginosa
VIM-7
VIM-28 P.aeruginosa
VIM-38 P.aeruginosa
VIM-5 P.aeruginosa
VIM-49 P.aeruginosa
VIM-19 E.coli
0.75
VIM-54 S.marcescens
VIM-27 K.pneumoniae
VIM-37 P.aeruginosa
VIM-40 E.cloacae
VIM-4 P.putida
VIM-43 P.aeruginosa
VIM-4 P.aeruginosa
VIM-1 clade
VIM-42 K.pneumoniae
VIM-29 E.coli
VIM-52 K.pneumoniae
VIM-26 K.pneumoniae
VIM-39 K.pneumoniae
VIM-33 K.pneumoniae
VIM-34 K.pneumoniae
Tree scale: 0.01
VIM-1 E.cloacae
VIM-35 K.oxytoca
VIM-32 K.oxytoca
VIM-1 K.pneumoniae
VIM-47 P.aeruginosa
VIM-13 P.aeruginosa
VIM-13 clade
0.99
VIM-30 P.aeruginosa
VIM-2e P.aeruginosa
VIM-17 P.aeruginosa
1.00
VIM-2b P.aeruginosa
VIM-10 P.aeruginosa
VIM-18 P.aeruginosa
VIM-36 P.aeruginosa
VIM-23 C.freundii
VIM-24 K.pneumoniae
VIM-50 P.aeruginosa
VIM-11 A.baumannii
VIM-3 P.aeruginosa
VIM-6 A.baumannii
VIM-31 E.cloacae
VIM-20 P.aeruginosa
VIM-2 clade
VIM-44 P.aeruginosa
VIM-15 P.aeruginosa
VIM-2d P.aeruginosa
VIM-14 P.aeruginosa
VIM-45 P.aeruginosa
VIM-2 P.aeruginosa
VIM-8 P.aeruginosa
VIM-9 P.aeruginosa
VIM-46 P.aeruginosa
VIM-41 P.aeruginosa
VIM-16 P.aeruginosa
VIM-51 K.pneumoniae
VIM-2c P.aeruginosa
VIM-2 P.putida
VIM-2 P.pseudoalcaligenes

**Supplementary Figure 6. Maximum likelihood phylogenetic tree of wtVIM-2 and 55 VIM-type variants.** The leaves in the tree indicate individual VIM-type variants labelled by number and host organism, with the wtVIM-2 sequence highlighted in blue. Other VIM-2 sequences are labelled in orange, and VIM-2 sequences that share the same host as wtVIM-2 have been labelled VIM-2b/c/d/e for disambiguation. The number beneath the branches indicate the normalized bootstrap value of 100 bootstrap replicates, and only bootstrap values above 0.75 are displayed. The major clades are noted by the labels on the right side.  The tree was generated using MEGA7.

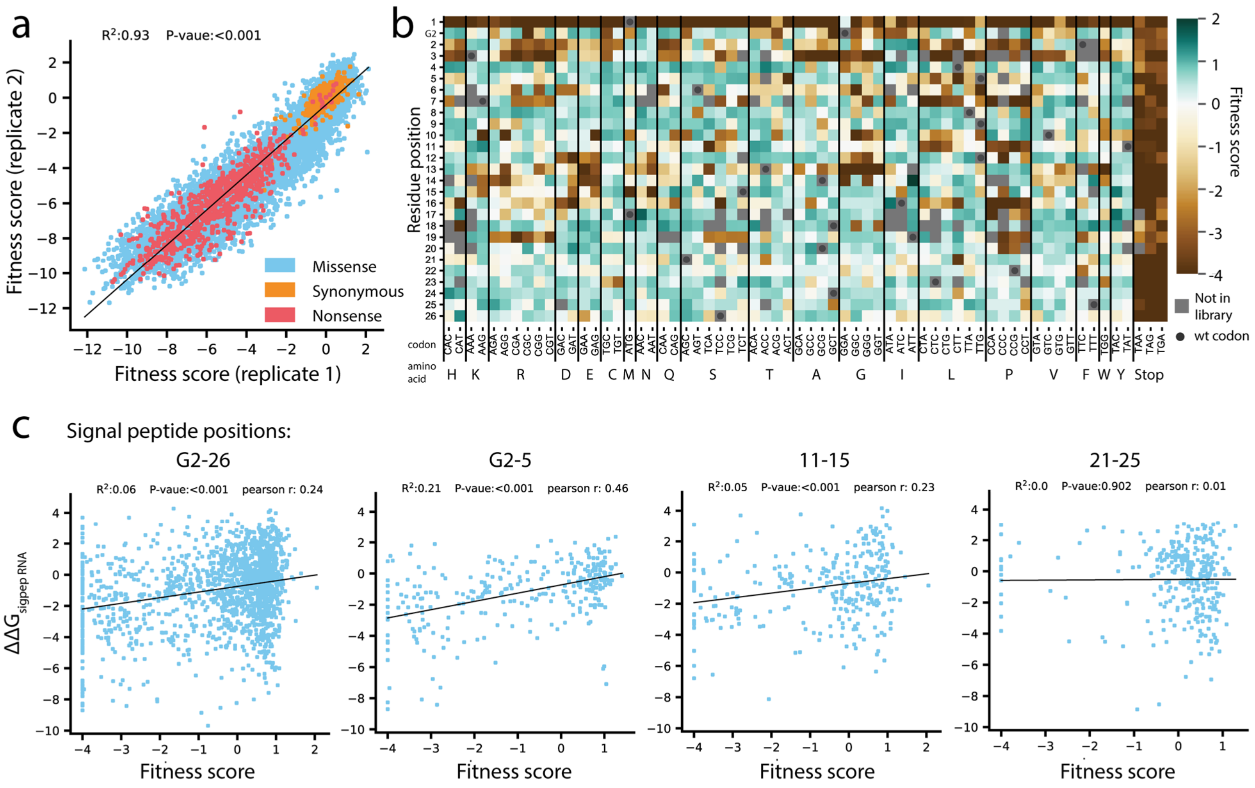


**Supplementary Figure 7. Codon variant fitness for VIM-2 under selection with 128 µg/mL AMP.** (**a**) Replicate correlation for fitness scores of all codon variants, with data points colored according to the type of mutation. The black line indicates the line of best fit for a linear regression between the replicates, with the $R^2$ and P-value displayed at the top. (**b**) Heatmap of fitness scores for the signal peptide of VIM-2. The residue positions are indicated on the y-axis (G2 refers to the Glycine 2 in the inhouse sequence that is not a part of wtVIM-2), while the codon mutations are indicated on the x-axis. (**c**) Plots showing correlation between the fitness score and predicted $\Delta\Delta G$ of RNA folding (energy contributions of the 5' UTR and the signal peptide region) for codon variants in the signal peptide. The $\Delta\Delta G$ is normalized to a multiple of the thermal energy factor $kT$, such that a $\Delta\Delta G$ of 1 indicates a $1 \times kT$ increase in $\Delta G$ of the variant over the wt. The label above each plot indicates the positions where codon variants were considered for the correlation, labelled as in panel (**b**.

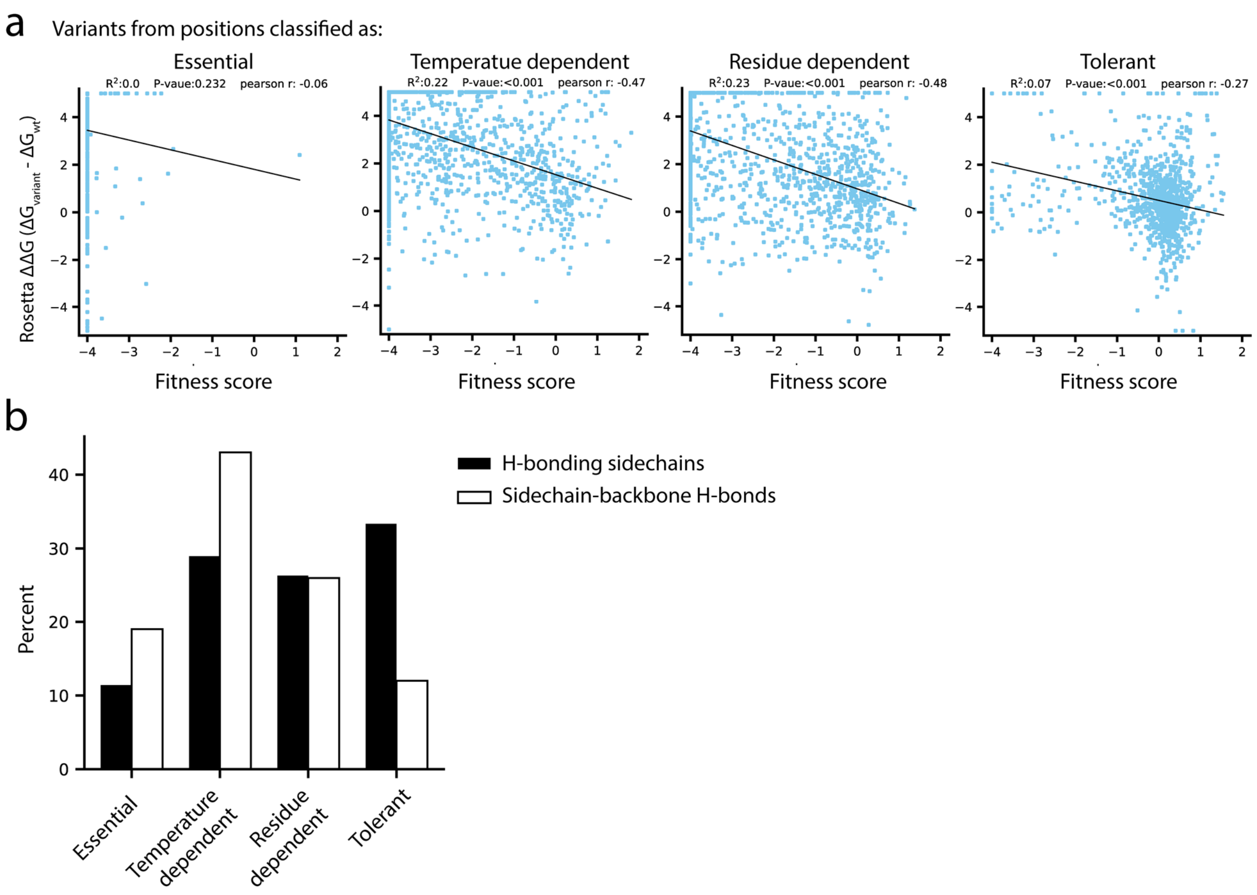


**Supplementary Figure 8. Rosetta Δ$\Delta G$ and hydrogen bonding behavior in relation to temperature dependence classifications.** Only positions in the catalytic domain (positions 27-266) are analyzed in these plots. (**a**) In each plot, Rosetta $\Delta\Delta G$ is correlated with fitness score for amino acid variants from the positions with the temperature dependence classification indicated at the top. (**b**) A comparison of proportion of h-bonding sidechains to the proportion of sidechain-backbone h-bonds within each temperature dependent classification. The black bars display the percentage of h-bonding sidechains within the wtVIM-2 catalytic domain (114 total) that fall within each temperature dependence classification. The white bars show the percentage of sidechain-backbone h-bonds (73 total) that are formed through sidechains of residues with the classification.

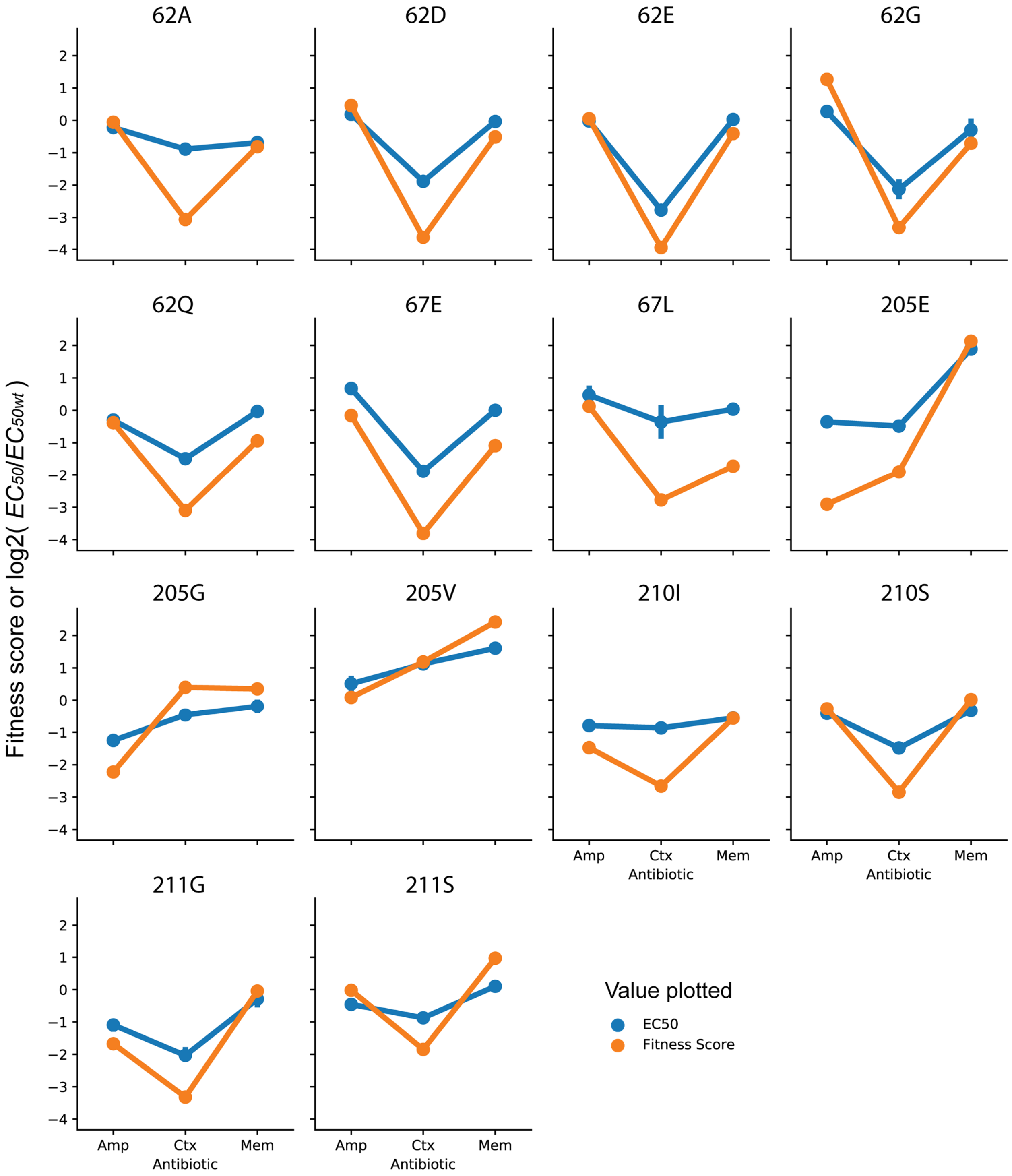
62A
62D
62E
62G
62Q
67E
67L
205E
205G
205V
210I
210S
211G
211S
Fitness score or log2( $EC_{50}/EC_{50wt}$ )
Amp
Ctx
Mem
Antibiotic
Value plotted
EC50
Fitness Score

**Supplementary Figure 9. Comparison of fitness scores and $EC_{50}$ of specificity variants.** Each plot shows the fitness score obtained from DMS (128 µg/mL AMP, 4.0 µg/mL CTX and 0.031 µg/mL MEM) or the $EC_{50}$ fitted from a dose response curve for the variant indicated. Fitness scores are plotted directly, while $EC_{50}$ values have been normalized to $\log_2(EC_{50}/EC_{50\ \mathrm{wt}})$ to allow them to be compared on the same scale as the fitness scores. $EC_{50}$ values of the same amino acid variants (different codon variants or same codon variants isolated and measured multiple times) show the average $EC_{50}$ with error bars indicating the 95% confidence interval.

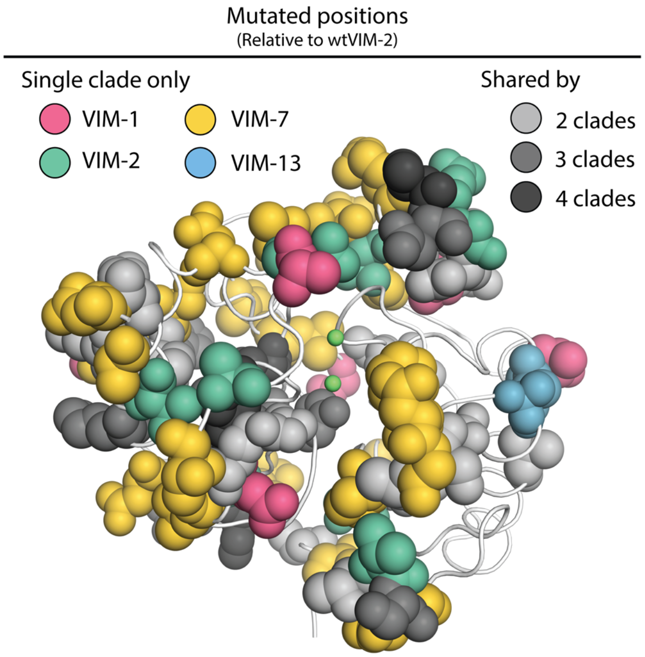


**Supplementary Figure 10. Mutated positions in natural VIM variants.** VIM-2 residues that are mutated in any natural variant are shown in sphere representation on the VIM-2 crystal structure (PDB: 5yd7), with positions mutated only within a single clade colored by clade and positions mutated in multiple clades colored by the number of clades.

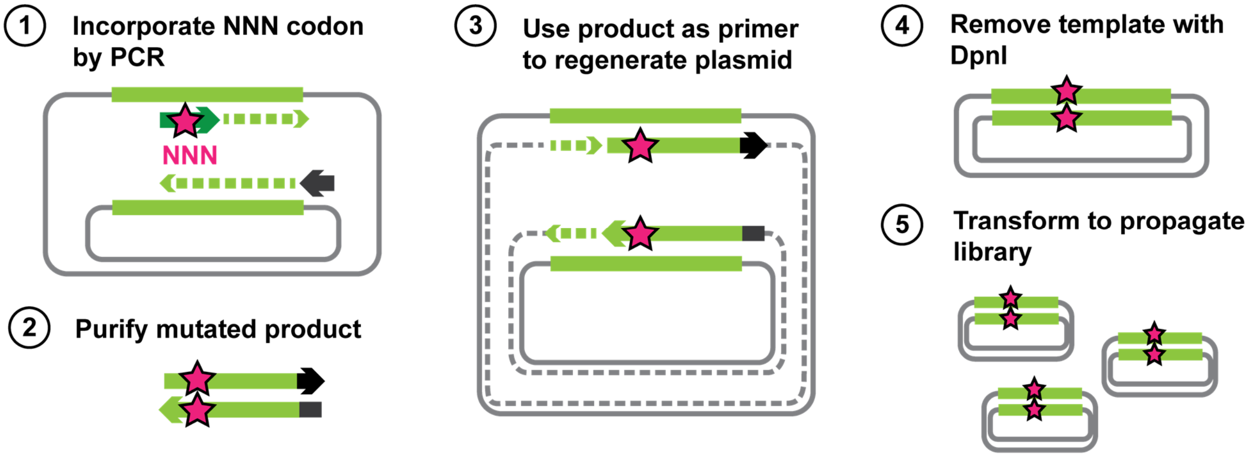


**Supplementary Figure 11. Outline of steps for generating all single amino acid variants of VIM-2.** The wtVIM-2 gene is encoded on a plasmid, and the first step is to extract the gene using a PCR. The forward primers carry an "NNN" mutation that is in frame with a specific codon of the wtVIM-2 gene, and all products will have a randomly mutated codon at that position. The mutated products are purified and used as primers to extend the rest of the plasmid using the same wtVIM-2 plasmid as the template, giving rise to the mutant VIM-2 plasmid library; the plasmid has a nick in both strands at the 5' end, but can anneal to each other to become circular. As a final clean up step, the second PCR products are digested with DpnI for 1 hour at 37°C to remove the wt template. The purified library is then transformed into E. coli for propagation and subsequently purified.

a

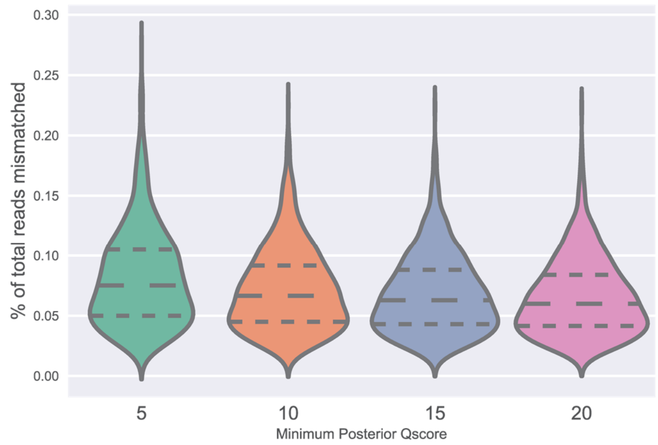


| | 5 | 10 | 15 | 20 |
|---|---|---|---|---|
| N | 804 | 804 | 804 | 804 |
| Mean | 0.0816 | 0.0719 | 0.0681 | 0.0653 |
| S.D. | 0.0408 | 0.0346 | 0.0327 | 0.0316 |

b

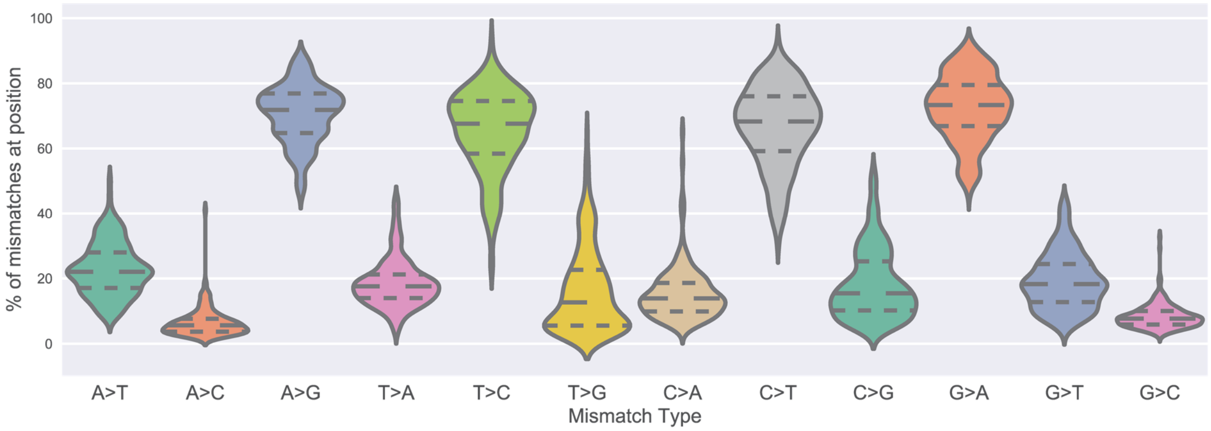


| | A>T | A>C | A>G | T>A | T>C | T>G | C>A | C>T | C>G | G>A | G>T | G>C |
|---|---|---|---|---|---|---|---|---|---|---|---|---|
| N | 165 | 165 | 165 | 201 | 201 | 201 | 206 | 206 | 206 | 232 | 232 | 232 |
| Mean | 22.8 | 6.48 | 70.72 | 18.52 | 65.32 | 16.16 | 15.23 | 66.63 | 18.14 | 72.51 | 19.18 | 8.31 |
| S.D. | 8.15 | 4.4 | 8.83 | 7.43 | 12.44 | 12.98 | 8.11 | 12.3 | 10.71 | 10.1 | 8.66 | 3.74 |
| % of Total | 5.57 | 1.71 | 18.32 | 5.12 | 19.45 | 5.04 | 3.14 | 15.53 | 3.92 | 16.75 | 3.77 | 1.67 |

**Supplementary Figure 12. Measurement of deep sequencing error rates using sequencing data of wtVIM-2 DNA.** (**a**) Violin plots showing the distribution of positional error rates observed in wtVIM-2 reads that pass filtering by having less than 20 mismatches between the forward and reverse reads and a minimum posterior Quality (Phred) score (5, 10, 15 or 20) at every read position. The positional error rate is calculated for every nucleotide position (804 positions in the VIM-2 gene) as the number of mutations divided by the total number of reads. The number of observations, distribution mean and standard deviation are listed below the x-axis. (**b**) Violin plots showing the distribution of proportions of specific mutations occurring due to sequencing error in the wtVIM-2 reads given a certain starting nucleotide. The proportions are calculated by dividing the number of a specific mutation at each wtVIM-2 nucleotide position (A>G, A>T, A>C) by the total number of mutations observed at that position. The proportions mutations arising from the same starting nucleotide sum to 100%. The number of observations, distribution mean, standard deviation and percentage of all observed mutations are listed below the x-axis.

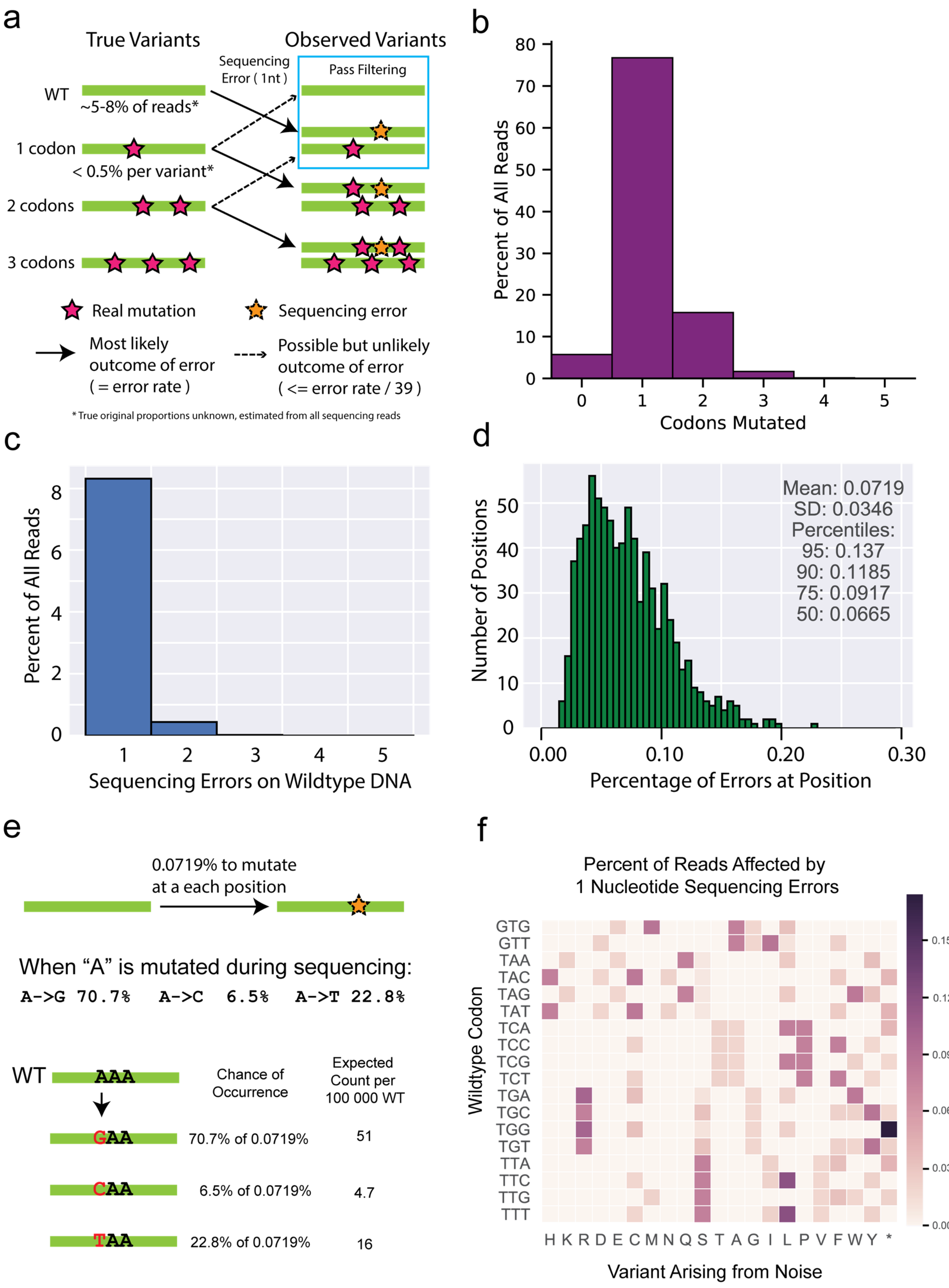
a
True Variants
Observed Variants
Sequencing Error ( 1nt )
Pass Filtering
WT
~5-8% of reads*
1 codon
< 0.5% per variant*
2 codons
3 codons
Real mutation
Sequencing error
Most likely outcome of error ( = error rate )
Possible but unlikely outcome of error ( <= error rate / 39 )
* True original proportions unknown, estimated from all sequencing reads
b
Percent of All Reads
Codons Mutated
c
Percent of All Reads
Sequencing Errors on Wildtype DNA
d
Mean: 0.0719
SD: 0.0346
Percentiles:
95: 0.137
90: 0.1185
75: 0.0917
50: 0.0665
Number of Positions
Percentage of Errors at Position
e
0.0719% to mutate at a each position
When "A" is mutated during sequencing:
A->G 70.7% A->C 6.5% A->T 22.8%
WT AAA
Chance of Occurrence
Expected Count per 100 000 WT
GAA 70.7% of 0.0719% 51
CAA 6.5% of 0.0719% 4.7
TAA 22.8% of 0.0719% 16
f
Percent of Reads Affected by 1 Nucleotide Sequencing Errors
Wildtype Codon
GTG GTT TAA TAC TAG TAT TCA TCC TCG TCT TGA TGC TGG TGT TTA TTC TTG TTT
H K R D E C M N Q S T A G I L P V F W Y *
Variant Arising from Noise

**Supplementary Figure 13. Rationale and support for estimating deep sequencing noise for filtering variants observed in non-selected libraries.** (**a**) The diagram depicts the main flow of variation when sequencing error is added on top of the existing variation. Sequences that are completely wt for every nucleotide occupy a sizeable portion of every library sequenced, and are also capable of being converted to any single amino acid variant through sequencing errors. Variants that already have a mutation are more likely to be mutated in a second codon and are likely to be filtered out. Additionally, single codon variants are already observed at fairly low frequencies, and are expected to contribute a negligible proportion of noise to any other variant. (**b**) A representative distribution of codon mutations in one of the selected libraries found after sequencing. In most cases at least 5% of the library is completely wt, indicated by those without any codon mutations. (**c**) A representative distribution of the proportion of reads that have a given number of sequencing errors observed when sequencing wtVIM-2 DNA. The reads with 0 sequencing errors are not shown to allow the smaller values to be resolved more clearly. (**d**) Distribution of sequence errors per position for every position in sequenced wtVIM-2 DNA after forward and reverse sequences have been merged and filtered by quality. This is the same distribution as the violin plot in **Supplementary Fig. 12a** with a minimum posterior quality score cutoff of 10. (**e**) An example calculation of expected noise arising from sequencing errors. (**f**) Visualization of the calculation shown in **e**) when applied across an entire all wt codons. The expected chance for each wt codon to become any of the other 9 codons adjacent by a single nucleotide substitution is first calculated and then aggregated at the amino acid level. Only a subset of the codons are shown as an example.

**Supplementary Table 1. Variants observed in each library group**

| | | | 37°C[a] | | 25°C[a] | |
|---|---|---|---|---|---|---|
| Group | Number of positions | Total possible variants | Both replicates[b] | At least one replicate[c] | Both replicates | At least one replicate |
| 1 | 39 | 819 | 808 | 811 | 808 | 812 |
| 2 | 39 | 819 | 812 | 813 | 811 | 814 |
| 3 | 39 | 819 | 803 | 807 | 802 | 807 |
| 4 | 39 | 819 | 802 | 813 | 809 | 812 |
| 5 | 39 | 819 | 801 | 812 | 804 | 811 |
| 6 | 39 | 819 | 775 | 793 | 789 | 796 |
| 7 | 33 | 693 | 682 | 686 | 678 | 690 |
| Total | 267 | 5607 | 5483 | 5535 | 5501 | 5542 |
| % coverage | | | 97.8% | 98.7% | 98.1% | 98.8% |

a) Non-selected libraries were grown, sequenced and filtered separately at 37°C and 25°C.
b) The observed number of variants that passed noise filtering in both sequencing replicates of the non-selected library.
c) The observed number of variants that passed noise filtering in at least one sequencing replicate of the non-selected library.

**Supplementary Table 2. Linear model output for DMS fitness scores under 4.0 μg/mL CTX selection**

| Predictor[a] | | Estimated effect[b] | Adjusted P-value ($\alpha$=0.05)[c] | Variance Explained[d] |
|---|---|---|---|---|
| (Intercept)[e] | | -1.40 | < 0.001 | |
| ASA[f] | | 2.75 | < 0.001 | 21% |
| Rosetta $\Delta\Delta G$[g] | | -0.27 | < 0.001 | 17% |
| Starting(WT) residue | C | -1.81 | < 0.001 | 10% |
| | D | -0.98 | < 0.001 | |
| | E | -0.58 | < 0.001 | |
| | F | -0.61 | < 0.001 | |
| | G | -0.65 | < 0.001 | |
| | H | -1.11 | < 0.001 | |
| | I | -0.40 | < 0.001 | |
| | L | -0.43 | < 0.001 | |
| | N | -0.96 | < 0.001 | |
| | S | -0.27 | 0.003 | |
| | V | -0.44 | < 0.001 | |
| | W | -1.54 | < 0.001 | |
| Variant residue | C | -1.49 | < 0.001 | 5% |
| | D | -0.46 | < 0.001 | |
| | E | -0.40 | 0.001 | |
| | G | -0.32 | 0.012 | |
| | P | -0.55 | < 0.001 | |
| | W | -0.43 | < 0.001 | |

a) Each predictor indicates a class of wtVIM-2 derived values that were used as explanatory variables to model a linear relationship with the observed fitness score.
b) The estimated effect is the predicted change in fitness score away from the intercept with a 1 unit increase in a continuous predictor or a binary change in a categorical predictor.
c) P-values indicates whether a predictor makes a significant contribution to the change fitness score, and are adjusted for a false discovery rate of 5% using the Benjamini-Hochberg procedure.
d) The adjusted $R^2$ of each predictor when correlated with fitness, which is a measure of how much variation in the fitness score can be explained by each predictor in the linear model.
e) The intercept is the average fitness of all variants where the continuous variable is 0 (ASA and Rosetta $\Delta\Delta G$) and the wt or variant residue is Ala.
f) ASA ranges from 0.0 to 1.0.
g) Rosetta $\Delta\Delta G$ ranges from -5.0 to 5.0 Rosetta energy units

**Supplementary Table 3. Linear model output for DMS fitness scores under 0.031 µg/mL MEM selection**

| **Predictor[a]** | | **Estimated effect[b]** | **Adjusted P-value (α=0.05)[c]** | **Variance Explained[d]** |
|---|---|---|---|---|
| (Intercept)[e] | | -1.30 | < 0.001 | |
| ASA[f] | | 2.64 | < 0.001 | 21% |
| Rosetta ΔΔ*G*[g] | | -0.30 | < 0.001 | 20% |
| Starting(WT) residue | C | -1.80 | < 0.001 | 10% |
| | D | -1.01 | < 0.001 | |
| | E | -0.61 | < 0.001 | |
| | G | -0.78 | < 0.001 | |
| | H | -1.20 | < 0.001 | |
| | I | -0.48 | < 0.001 | |
| | L | -0.52 | < 0.001 | |
| | N | -0.59 | < 0.001 | |
| | Q | -0.32 | 0.025 | |
| | R | -0.28 | 0.007 | |
| | S | -0.21 | 0.023 | |
| | V | -0.41 | < 0.001 | |
| | W | -1.52 | < 0.001 | |
| Variant residue | C | -1.60 | < 0.001 | 5% |
| | D | -0.47 | < 0.001 | |
| | E | -0.34 | 0.006 | |
| | G | -0.33 | 0.009 | |
| | P | -0.52 | < 0.001 | |
| | W | -0.42 | 0.001 | |

a) Each predictor indicates a class of wtVIM-2 derived values that were used as explanatory variables to model a linear relationship with the observed fitness score.
b) The estimated effect is the predicted change in fitness score away from the intercept with a 1 unit increase in a continuous predictor or a binary change in a categorical predictor.
c) P-values indicates whether a predictor makes a significant contribution to the change fitness score, and are adjusted for a false discovery rate of 5% using the Benjamini-Hochberg procedure.
d) The adjusted $R^2$ of each predictor when correlated with fitness, which is a measure of how much variation in the fitness score can be explained by each predictor in the linear model.
e) The intercept is the average fitness of all variants where the continuous variable is 0 (ASA and Rosetta ΔΔ*G*) and the wt or variant residue is Ala.
f) ASA ranges from 0.0 to 1.0.
g) Rosetta ΔΔ*G* ranges from -5.0 to 5.0 Rosetta energy units

**Supplementary Table 4. Inferred specificity of residues in wtVIM-2**

| Specificity | Positions |
|---|---|
| AMP | 39, 55, 57, 63, 117, 119, 142, 143, 153, 196, 219, 222, 243, 248 |
| CTX | 62, 67, 68 |
| AMP or CTX | 60, 61, 202, 205, 210, 211, 216 |
| MEM | 201 |

**Supplementary Table 5. Natural VIM variants from CARD and NCBI**

| Accession[a] | Name | Host organism | Mutations vs wtVIM-2 | Vim clade |
|---|---|---|---|---|
| tr\|A0A023UGS6 | VIM-1 | *Klebsiella pneumoniae* | 26 | VIM-1 |
| tr\|Q2HQ50 | VIM-1 | *Enterobacter cloacae* | 28 | VIM-1 |
| tr\|Q6B3N4 | VIM-2 | *Pseudomonas pseudoalcaligenes* | 1 | VIM-2 |
| tr\|A4GRB6[b] | VIM-2 | *Pseudomonas aeruginosa* | - | VIM-2 |
| tr\|A0A0M3THU2 | VIM-2 | *Pseudomonas putida* | 2 | VIM-2 |
| tr\|D1MEN9 | VIM-2b | *Pseudomonas aeruginosa* | 2 | VIM-2 |
| tr\|F8S2I1 | VIM-2c | *Pseudomonas aeruginosa* | 2 | VIM-2 |
| tr\|A0A0M4FPL5 | VIM-2d | *Pseudomonas aeruginosa* | 2 | VIM-2 |
| tr\|B8QIQ9 | VIM-2e | *Pseudomonas aeruginosa* | 2 | VIM-2 |
| gb\|AAG27703.1 | VIM-3 | *Pseudomonas aeruginosa* | 3 | VIM-2 |
| gb\|CAJ32502.1 | VIM-4 | *Pseudomonas aeruginosa* | 25 | VIM-1 |
| tr\|B3WFR2 | VIM-4 | *Pseudomonas putida* | 26 | VIM-1 |
| tr\|Q6SKX7 | VIM-5 | *Pseudomonas aeruginosa* | 28 | VIM-1 |
| tr\|A7L375 | VIM-6 | *Acinetobacter baumannii* | 3 | VIM-2 |
| gb\|CAD61201.1 | VIM-7 | *Pseudomonas aeruginosa* | 70 | VIM-7 |
| gb\|AAS13759.1 | VIM-8 | *Pseudomonas aeruginosa* | 2 | VIM-2 |
| gb\|AAS13760.1 | VIM-9 | *Pseudomonas aeruginosa* | 2 | VIM-2 |
| gb\|AAS13761.1 | VIM-10 | *Pseudomonas aeruginosa* | 2 | VIM-2 |
| tr\|Q0GN98 | VIM-11 | *Acinetobacter baumannii* | 2 | VIM-2 |
| tr\|Q2HY42 | VIM-13 | *Pseudomonas aeruginosa* | 33 | VIM-13 |
| tr\|A0SWU7 | VIM-14 | *Pseudomonas aeruginosa* | 2 | VIM-2 |
| tr\|B4YAJ6 | VIM-15 | *Pseudomonas aeruginosa* | 2 | VIM-2 |
| tr\|B4YAJ7 | VIM-16 | *Pseudomonas aeruginosa* | 2 | VIM-2 |
| tr\|B5KVR9 | VIM-17 | *Pseudomonas aeruginosa* | 2 | VIM-2 |
| gb\|CAO83029.1 | VIM-18 | *Pseudomonas aeruginosa* | 6 | VIM-2 |
| gb\|ACY29468.1 | VIM-19 | *Escherichia coli* | 25 | VIM-1 |
| gb\|ACV13198.1 | VIM-20 | *Pseudomonas aeruginosa* | 2 | VIM-2 |
| tr\|A0A1B4XBF6 | VIM-23 | *Citrobacter freundii* | 2 | VIM-2 |
| tr\|E0AFT6 | VIM-24 | *Klebsiella pneumoniae* | 2 | VIM-2 |
| tr\|E5BDC6 | VIM-26 | *Klebsiella pneumoniae* | 26 | VIM-1 |
| tr\|F6L7B6 | VIM-27 | *Klebsiella pneumoniae* | 27 | VIM-1 |
| tr\|F8UTU0 | VIM-28 | *Pseudomonas aeruginosa* | 25 | VIM-1 |
| gb\|AFP99885.1 | VIM-29 | *Escherichia coli* | 27 | VIM-1 |
| gb\|AET05999.1 | VIM-30 | *Pseudomonas aeruginosa* | 2 | VIM-2 |
| tr\|I3RJZ3 | VIM-31 | *Enterobacter cloacae* | 3 | VIM-2 |
| tr\|H6UQF5 | VIM-32 | *Klebsiella oxytoca* | 27 | VIM-1 |
| gb\|AFP99175.1 | VIM-33 | *Klebsiella pneumoniae* | 27 | VIM-1 |
| gb\|AFN88953.1 | VIM-34 | *Klebsiella pneumoniae* | 27 | VIM-1 |
| gb\|AGC50805.1 | VIM-35 | *Klebsiella oxytoca* | 27 | VIM-1 |
| tr\|L7US14 | VIM-36 | *Pseudomonas aeruginosa* | 2 | VIM-2 |
| gb\|AGC50807.1 | VIM-37 | *Pseudomonas aeruginosa* | 26 | VIM-1 |
| tr\|M1JB36 | VIM-38 | *Pseudomonas aeruginosa* | 26 | VIM-1 |
| tr\|A0A059PYQ6 | VIM-39 | *Klebsiella pneumoniae* | 27 | VIM-1 |
| tr\|X5JRU4 | VIM-40 | *Enterobacter cloacae* | 26 | VIM-1 |
| tr\|A0A0K0PWY0 | VIM-41 | *Pseudomonas aeruginosa* | 2 | VIM-2 |
| tr\|A0A0C5GGI4 | VIM-42 | *Klebsiella pneumoniae* | 25 | VIM-1 |
| gb\|AJP67511.1 | VIM-43 | *Pseudomonas aeruginosa* | 26 | VIM-1 |
| tr\|A0A0H4J4X8 | VIM-44 | *Pseudomonas aeruginosa* | 2 | VIM-2 |
| tr\|A0A0H4IR82 | VIM-45 | *Pseudomonas aeruginosa* | 2 | VIM-2 |
| tr\|A0A0K0PX71 | VIM-46 | *Pseudomonas aeruginosa* | 3 | VIM-2 |
| tr\|A0A0P0M6J0 | VIM-47 | *Pseudomonas aeruginosa* | 32 | VIM-13 |

| | | | | |
|---|---|---|---|---|
| tr\|A0A125R6L1 | VIM-49 | *Pseudomonas aeruginosa* | 29 | VIM-1 |
| tr\|A0A125R6L2 | VIM-50 | *Pseudomonas aeruginosa* | 3 | VIM-2 |
| tr\|A0A140EA28 | VIM-51 | *Klebsiella pneumoniae* | 2 | VIM-2 |
| tr\|A0A218KGB0 | VIM-52 | *Klebsiella pneumoniae* | 26 | VIM-1 |
| tr\|A0A1P8SAK1 | VIM-54 | *Serratia marcescens* | 26 | VIM-1 |

a. Accessions starting with 'gb' are from Genbank, while those starting with 'tr' are from TrEMBL
b. The same sequence as wtVIM-2 in this study.
